# Embedding bioprinting of low viscous, photopolymerizable blood-based bioinks in a self-healing transparent supporting bath

**DOI:** 10.1101/2024.05.29.596452

**Authors:** Monize Caiado Decarli, Helena P. Ferreira, Rita Sobreiro-Almeida, Filipa C. Teixeira, Tiago R. Correia, Joanna Babilotte, Jos Olijve, Catarina A. Custódio, Inês C. Gonçalves, Carlos Mota, João F. Mano, Lorenzo Moroni

## Abstract

Protein-based hydrogels have great potential to be used as bioinks for biofabrication-driven tissue regeneration strategies due to their innate bioactivity. Nevertheless, their use as bioinks in conventional 3D bioprinting is impaired due to their intrinsic low viscosity. Using embedding bioprinting, a liquid bioink is printed whithin a support that physically holds the patterned filament. Inspired by the recognized microencapsulation technique complex coacervation, we introduce crystal self-healing embedding bioprinting (CLADDING) based on a highly transparent crystal supporting bath. The suitability of distinct classes of gelatins was evaluated (i.e., molecular weight distribution, isoelectric point and ionic content), as well as the formation of gelatin-gum arabic microparticles as a function of pH, temperature, solvent and mass ratios. Characterizing and controlling this parametric window resulted in high yields of support bath with ideal self-healing properties for interaction with protein-based bioinks during bioprinting. This support bath achieved transparency, which boosted light permeation within the bath. CLADDING bioprinted constructs fully composed of platelet lysates encapsulating a co-culture of human mesenchymal stem cells and endothelial cells were obtained, demonstrating high-dense cellular network with excellent cell viability and stability over a month. CLADDING broadens the spectrum of photocrosslinkable materials with extremely low viscosity that can now be bioprinted with sensitive cells using embedding bioprinting without any additional support.

## Introduction

Several protein-based hydrogels have great potential to be used as bioinks for biofabrication-driven tissue regeneration strategies due to their innate bioactivity.^[1,2]^ Some hydrogel examples are based on collagen, fibronectin, peptide- and extracellular matrix-based inks, and blood derivatives, including from human origin.^[3]^ Two of such blood derivatives, platelet lysates (PL) and platelet-rich plasma (PRP) from platelet concentrates of whole blood, have been used as a source of autologous material for 3D culture platforms and tissue engineering with promising results.^[2,4–6]^ PL and PRP are great sources of highly biocompatible xeno-free biomaterials with intrinsic cell-recruiting and pro-regenerative capacity.^[7]^ PL consists of a natural repair cocktail mainly composed of albumin (41%), bioactive molecules (e.g. growth and pro-angiogenic factors and cytokines), structural proteins (e.g. fibrinogen, fibronectin, and vitronectin), as well as antimicrobial peptides and lipid-binding molecules, among others.^[2,8]^

Like other hydrogels with intrinsic high fluidity and low viscosity, PL and PRP cannot be bioprinted by itself despite their remarkable biological properties. It has also been shown that living cells cannot be loaded on PL and PRP without prior modification because they can easily sink to the bottom of the material, which hinders their use for biofabrication applications.^[9]^ Recently, extensive research has been conducted to overcome this limitation by combining PL and PRP with several additives to increase their viscosities and make them printable. Alternatively, structures made with PL and PRP have been obtained via molding and casting (**Supplementary Table 1**).^[4,9–18]^ So far, the bioprinting of only PL or other blood derived low viscosity bioinks without any external thickener or support mold has not been reported.

Nevertheless, it is possible to extrude these low viscous bioinks without addition of any thickener by the use of embedded bioprinting. In this technique, a bioink is printed within a hydrogel support that physically holds the patterned printed filament.^[19]^ Several approaches have been already efficiently established to prevent structural collapse, including pyrogenic silica,^[20]^ synthetic agranular gels made of Carbopol,^[21]^ pluronic F-127,^[19]^ and silicon.^[22]^ Support baths have also been described using biopolymers such as gelatin,^[23]^ hyaluronic acid,^[24]^ xantham-gum,^[25]^ agarose,^[26]^ and microgels with ECM molecules^[27]^, which exhibited interesting self-healing and viscoelastic responses. Moreover, since some of these biopolymers are thermosensitives, they can be easily removed by melting in a cell incubator, thus being sacrificial support baths.

Using the freeform reversible embedding of suspended hydrogels (FRESH,V.2), a supporting bath was also obtained with gelatin–gum arabic using coacervation.^[28]^ Coacervation is one of the oldest and possibly the most widely recognized microencapsulation techniques.^[29]^ Simple coacervation is performed with non-ionized groups and usually with one colloid, whereas complex coacervates deal with two or more oppositely charged macromolecules.^[29,30]^ The gelatin–gum arabic pair has been used in many applications as a model protein-polysaccharide for complex coacervation since 1920.^[30]^ Comparing gelatin coacervation to a range of materials for obtaining support baths, Shiwarski et al. made a compilation of many biological materials that were successfully obtained by embedding printing.^[31]^ Using these studies as basis, we asked whether coacervation with gelatin-gum arabic could be enhanced in self-healing and transparency capability to allow bioprinting of low viscous photocroslinkable blood derivatives in mild conditions for bioprinting sensitive tissues containing endothelial and stem cells. As a result, we present herein a method to generate a crystal self-healing embedding bioprinting (CLADDING). CLADDING reflects our application of materials over other materials to obtaining resistance and insulation in a lightweight approach, analogous to the use of cladding composites in the construction field.

To develop CLADDING, we investigated how gelatin, which is a molecule from natural origin that can vary depending on the animal source, extraction procedure and composition,^[32]^ affects the generation of efficient coacervates with gum arabic (**Figure 1A**). We thus examined different types of gelatins with defined gel strength (50, 100, 150, 200, 250) and a non-defined gel strength (50–120), in terms of average molecular weight distribution (AMW), isoelectric point (IEP) and ionic content. Since coacervation is a colloidal phenomenon, the main parameters investigated herein were the appropriate solvent, nature/ratio of the colloid, temperature and pH. All these parameters play a major role in reducing the solubility of the colloid, making it precipitate and resulting in high yields of the coacervate phase.^[29,30]^ We also studied how the coacervate procedure impacts the final characteristics of the bioprinted scaffolds, such as printing fidelity and resolution (**Figure 1B**). The supporting baths were extensively characterized by rheology in terms of flow and self-healing behavior, as well as for achieving transparency to enhance photocrosslinking. Photocrosslinkable bovine serum albumin methacrylated (BSAMA), which is easier and cost-effective to obtain, and more uniform than PL,^[33]^ was used as proof-of-concept as a highly fluid and non-printable protein-based bionk. Based on BSAMA results, methacrylated platelet lysates (PLMA) were further explored (**Figure 1C**). After examinating properties and characteristics of PLMA and BSAMA photocrosslinkable hydrogels, PLMA bioink encapsulating human Mensenchymal Stromal Cells (hMSC) in single and co-cultures with human Umbilical Vein Endothelial Cells (hUVECs) were extruded to obtain bioprinted constructs in several shapes and sizes (**Figure 1D**). The visual aspect of the support bath obtained by CLADDING can be seen in **Figure 1E** and the optimized bioprinter set up for embedding bioprinting with simoutaneously photocrosslinking in **Figure 1F**. This study demonstrates the feasibility of bioprinting low viscousity blood components without thickners, which yielded bioprinted constructs with precise printing fidelity, high stability and cell viability.

**Figure 1:**
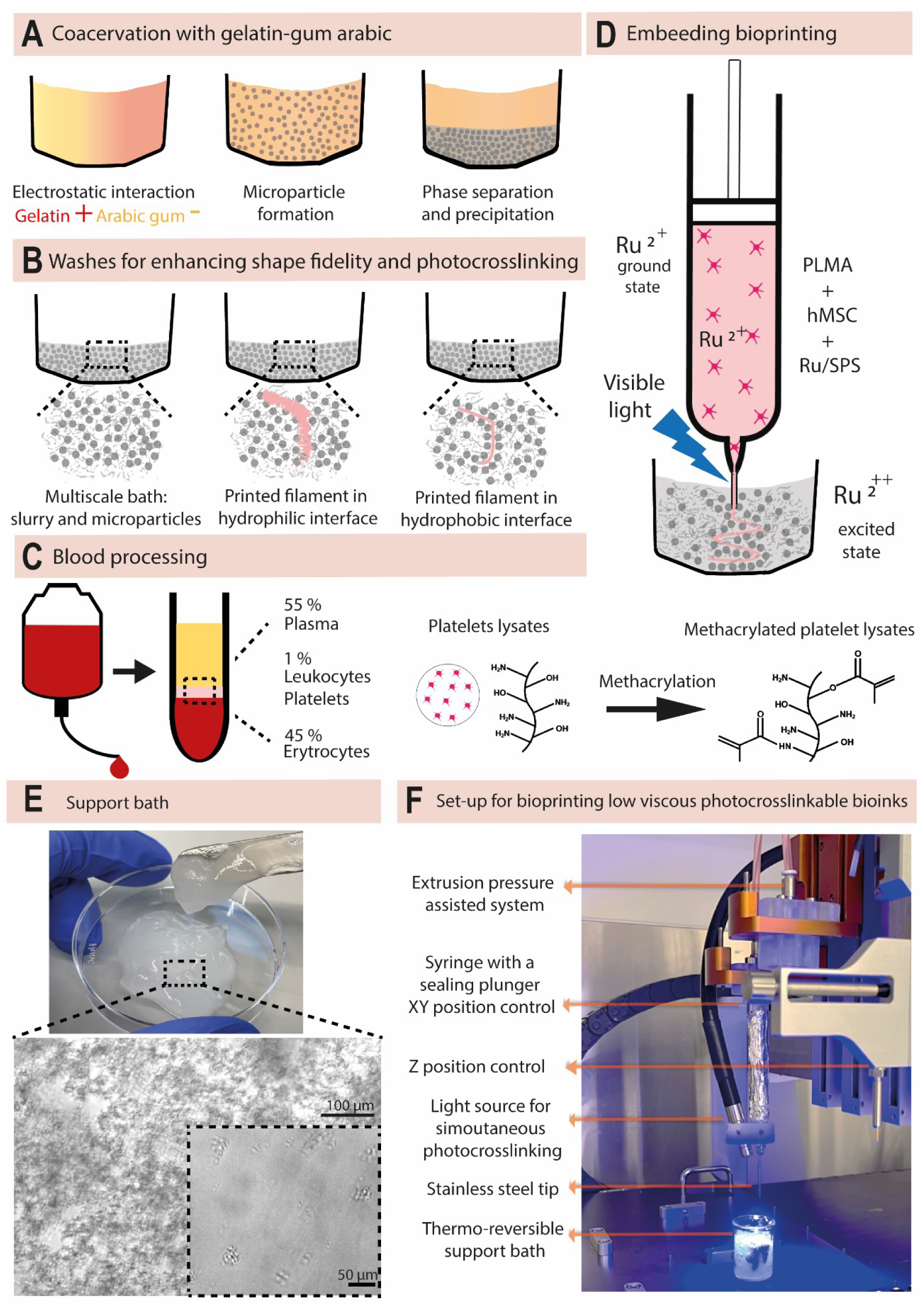
Schematic illustration for bioprinting low viscosity and photocrosslinkable bioinks by the CLADDING method. **A)** An established coacervation procedure involving two oppositely charged materials was intensively explored. Gelatin (positively charged when IEP is around 5) and gum arabic (negatively charged) formed a stable polyelectrolyte complex, followed by sequential phases of emulsification, phase separation, and precipitation. After complete precipitation, the coacervate phase was collected and stored. **B)** Micro-macro character of the support bath. Right before bioprinting, several cycles of washes and centrifugation using CaCl_2_ or Tween 20 were performed to compact the coacervate into a slurry, used as the support bath, and to increase its hydrophobic character, improving filament bioprinting fidelity (pink line representing the bioink). **C)** Platelets were obtained from human blood by the differentiation centrifugation process. PL solution was functionalized by addition of methacryloyl groups. Commercial BSA was acquired and methacrylated following the same procedure as for PLMA. The obtained BSAMA was used as a control. **D)** PLMA bioink (pink solution and printed filament) was prepared by mixing Ruthenium/sodium persulfate photoinitiator system (Ru^2+^, ground state) with hMSCs in single or co-cultures with hUVECs. Bioprinting procedure was performed along with a simultaneous photocrosslinking approach using visible light (Ru^2++^, excited state). **E)** Micro- and macroscopic views of the enhanced support bath. The formation of microparticles made of gelatin–gum arabic was observed using bright-field microscopy (scale bar = 50 µm), which showed a strong tendency to aggregate, forming a typical colloid (scale bar = 100 µm), thus making a multiscale support bath. **F)** The optimized set-up for controlling several parameters for bioprinting low viscosity, photocrosslinkable bioinks using embedding bioprinting can be seen.

## Results and Discussion

### Physico-chemical characterization of gelatin B

Aiming to further understand the physical phenomena behind coacervation to better control the bioprinting process, we investigated the properties and characteristics of the widely used component of supporting baths, gelatin, such as type, AMW, distribution of gelatin subunits, IEP, conductivity and ion content (calcium, sodium, magnesium, chlorides, and sulfates). Afterwards, we investigated the parameters that could affect the electrostatic interaction between oppositely charged gelatin (positive) and gum arabic (negative) pair, since this interaction is the major driving force for complex coacervation.

Gelatin can be obtained from bovine, pig or fish animal based by-products. Type A is obtained by acid hydrolysis and type B by alkaline hydrolysis. Thus, gelatin is a solubilized collagen derived from acid or lime treatment.^[32]^ Since coacervation is a process with high sensitivity to variation in pH and ionic strength, we first analyzed the properties of type A and B gelatins with the same gel strength and from the same supplier (**Supplementary** Figure 1). The gel strength for gelatin is commercially called “bloom”. The bloom value aims to correlate the physical and mechanical strength of gelatin-based products, such as capsules in the pharmaceutical industry to classify them as hard or soft capsules. The AMW of type A (136.6 kDa) was lower than type B (157.1 kDa), and gelatin type B presented a monodispersed and sharper distribution, ideal for obtaining homogeneous and reproducible coacervates. IEP is another crucial parameter, since it determines the charge balance between gelatin and gum arabic for achieving high precipitation yields; an ideal IEP for gelatin in this pair with gum arabic for coacervation is approximately 5. Gelatin type A showed an IEP of 8.71, a high charge unsuitable for coacervates with gum arabic, whereas type B had an ideal IEP equal to 5. Finally, conductivity was assessed, and gelatin type A showed a higher value than type B (130 and 88 µS/cm, respectively). Lower values in conductivity is preferable since higher ones can affect the coacervation. Together, these results strongly indicate that gelatin type B is preferred for further study.

Once gelatin type B was defined as the ideal type for this coacervation procedure, we investigated a set of gelatins with defined and non-defined properties. As defined ones, gel strengths of 50, 100, 150, 200, and 250 (named type “I”) within the normal range of 50–280 for commercial products were examined.^[32]^ As a matter of comparison, a non-defined gelatin with gel strength of 50-120 (named type “II”) was chosen. The main parameters of AMW, IEP, and conductivity of all these gelatins can be seen in **Table 1**.

**Table 1:** Average molecular weight, isoelectric point, and conductivity of a set of gelatins with defined and non-defined gel strengths.

| | Gelatin gel strength (bloom) | Average molecular weight (kDa) | Isoelectric point (IEP) | Conductivity ($\mu\text{S}/\text{cm}$) |
| --- | --- | --- | --- | --- |
| **Type B** | 50 | $89.62 \pm 2.6$ | 5.03 | 87 |
| | 100 | $62.55 \pm 2.6$ | 4.97 | 97 |
| | Non-defined 50-120 | $95.06 \pm 2.5$ | 5.63 | 290 |
| | 150 | $198.42 \pm 6.4$ | 5 | 93 |
| | 200 | $180.06 \pm 12.3$ | 5.04 | 64 |
| | 250 | $157.05 \pm 8.9$ | 5.02 | 88 |
| **Type A** | 250 | $136.64 \pm 2.8$ | 8.71 | 130 |

Analyzing the molecular weight distribution (**Figure 2A**), 250 bloom gelatin presented the most monodispersed pattern with the sharpest distribution, whereas 200 bloom had a similar but less pronounced pattern. For 50, 100 and 150 blooms and blend 50–120 gelatins (**Figure 2A**), a polydispersed pattern was observed. Similarly, in terms of the components of gelatin ranging from small subunits (0–70 kDa), α (70–115 kDa), β (115–210 kDa), γ (210–360 kDa) to larger ones such as microgels (>360 kDa), the 250 bloom gelatin showed roughtly even 20% across the sizes, whereas 150 and 200 blooms had a similar distribution of components, but less narrow distributed (**Figure 2B**). Conversely, lower blooms and 50–120 bloom blend were shown to be mainly composed of small subunits.

**Figure 2:**
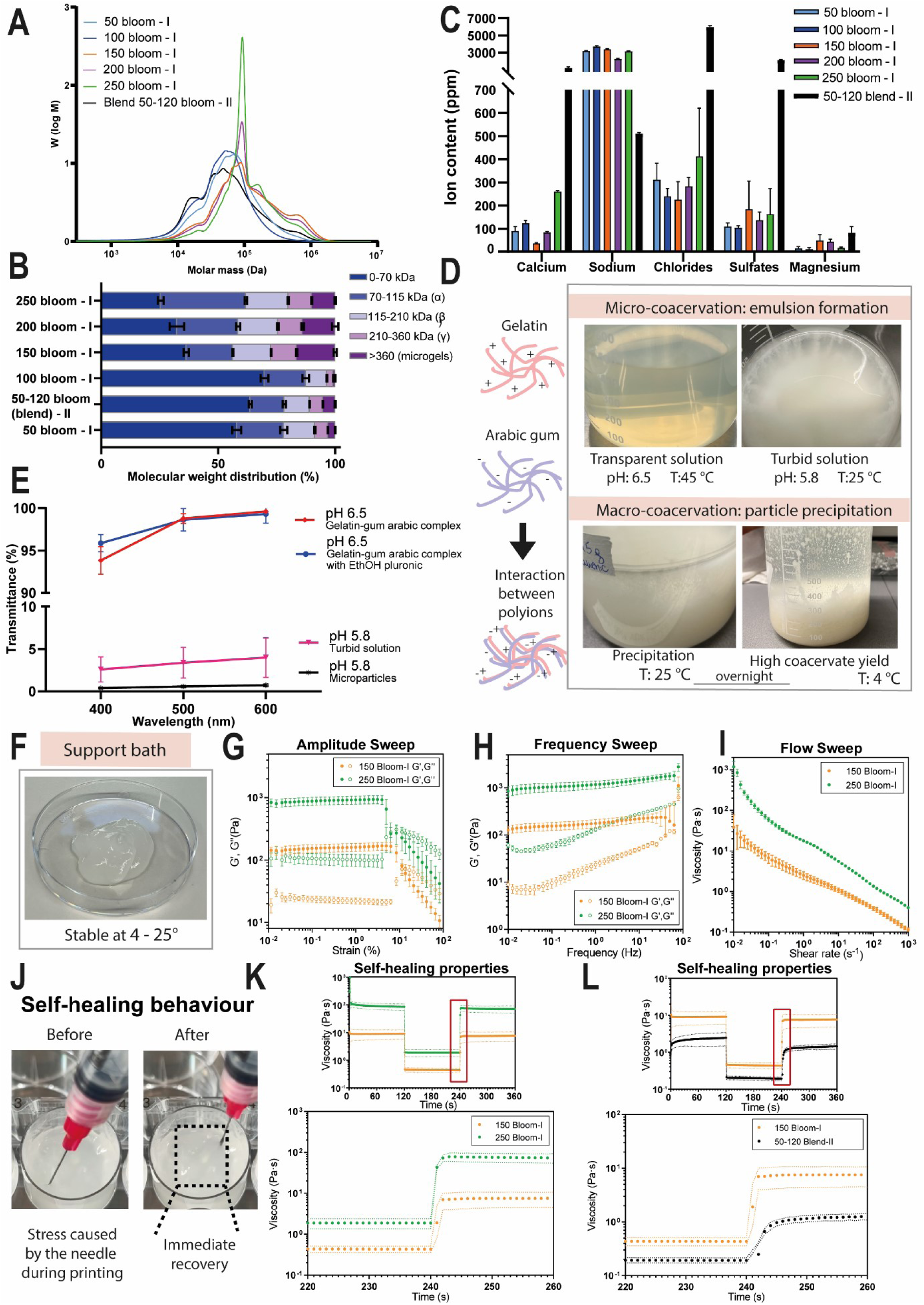
Properties and characteristics of gelatins type B with five defined gel strengths (bloom-I) with a non-defined gel strength (50-120 blend-II). Comparison in terms of **A)** molecular weight distribution; **B)** distribution of subunits based on size, and **C)** ion content. **D)** Micro-coacervation step initiated by lowering the pH and temperature of the system allowing emulsion formation.Macro-coacervation step intensified by a second dropping in temperature and incubation overnight resulted in a clear interface between the fluid phase and the coacervate, resulting in high yields of the coacervate phase. **E)** Quantitative characterization of the coacervate by measurements of turbidity as a function of pH to obtain the light thansmittance in all these steps. **F)** Gelatin-gum arabic coacervates developed by CLADDING ready for embedding bioprinting in temperatures ranging from 4 to 25 °C. **G–I**) Rheological characterization of supporting baths of gelatins with defined bloom 150 and 250, **(G)** including strain response**; (H)** frequency sweep and **(I)** flow behavior. **J)** Relevance of self-healing behavior for bioprinting and its assessment when comparing **(K, L).** Self-healing of supporting baths using **(K)** defined gelatins of 150 and 250 bloom compared to a **(L)** non defined bloom of 50-120 blend gelatin.

Since the major driving factor for coacervation is the electrostatic interaction, it was crucial to quantify the ion content in the gelatin since it might affect the whole ionic strength of the gelatin-gum arabic chains during the complexation process.^[30]^ The amounts of magnesium, calcium, chloride, sulfate, and sodium ions were determined in the whole set of gelatins considered in this study (**Figure 2C**). In gelatin I with defined gel strength, the ion content was similar, with a slightly increasing trend with increased gel strength. Magnesium was identified in small amounts (10–40 ppm), followed by calcium, chlorides, and sulfates which were identified in a range of 100–400 ppm. Sodium ions were greatest, measuring 2000– 4000 ppm. Such levels are due to the deonization step that gelatins with defined properties are often submitted to after extraction, using anionic/cationic resins to neutralize the unbalanced ionic content. After deionization, the pH was around 5, which is the same of IEP. Thus, to avoid a pH around the same value of IEP, which can compromise many applications such as coacervation procedures, an adjustment using NaOH is performed, explaining the high amount of observed sodium. Since the quantity of salt affects the ionic strength of the solution, previous reports have shown that when the concentration of NaCl is too low, coacervation may be suppressed^[34]^. Conversely, if the concentration is too high, the excess of chloride turns out to be the problem since it affects complex coacervation processes.^[35]^ Overall, we conclude that these values of salt concentration are the optimal ones for coacervation to take place with gum arabic and were identified in all the gelatins with defined properties.

The ion content in the gelatin with non-defined bloom 50-120 showed a different pattern. Magnesium, calcium, sulfate, and chloride were higher in blend 50-120 compared to the defined gelatins, at 2–3×, 10×, 10×, and 20× levels, respectively. Sodium was 4–5× lower in blend 50-120 compared to the defined gelatins. Sodium sulfate *per se* can be used as a precipitation agent during a simple coacervate process to separate gelatin.^[29,32]^ Thus, the high amount of sulfate and low amount of sodium in blend 50-120 can certainly disturb the electrostatic interaction with gum arabic and weaken the whole coacervation procedure. It is the first time that the ion content of gelatin is characterized as a function of gel strength and source. This characterization is of high relevance for effectively using gelatin in many biomedical applications.

Considering the analysis of the different gelatins, we prioritized monodispersity and homogeneity for molecular weight, as well as fixed IEP and low ion content for obtaining coacervates in a reproducible way. Using these criteria, we selected 150 and 250 bloom defined gelatins I for the coacervate procedure, and the non-defined blend 50-120 gelatin II as a control.

### Coacervation process to produce supporting baths: protein/polysaccharide ratio, solvent, pH and temperature

The CLADDING method is a coacervation procedure based on a three-phase system involving the solvent, the positively and negatively charged macromolecules. Herein, we divided this process into two main steps, called micro- and macro-coacervation. The micro-coacervation (**Figure 2D**) starts with a preparation of an aqueous solution to dissolve the gelatin (150, 250 and 50-120 blend) and gum arabic. Even though previous work have pointed the optimum mass ratio between gelatin and gum arabic as 1:1^[28,30]^ we identified that a clear biphasic system as a result of an efficient precipitation could only be seen with a ratio equal to 4:1 (**Supplementary** Figure 2). In terms of yield, 4:1 ratio produced two times more embeeding support (12.5 ± 0.2 g) than compared to 1:1 ratio (6.1 ± 0.4 g). In terms of waste, 4:1 ratio produced 40% less waste (7.8 ± 0.1 g) than compared to 1:1 ratio (12.8 ± 0.6 g) per 100 mL of solution. (**Supplementary Table 2**). Thus, the 4:1 gelatin-gum arabic makes the coacervation procedure more efficient.

The ideal condition for complete dissolution of these components, verified by a homogeneous and transparent solution, is in distilled water (dwater), pH 6.5, 45 °C in continuous vigorous stirring for 1 h (**Figure 2D**). After that, absolute ethanol in the same amount of water is added to the mixture, to achieve a final solvent mixture of 50% water/50% ethanol. Since the ethanol fraction contributes to further precipitation, we observed that if mixed in the previous step for dissolving gelatin and gum arabic as previously reported,^[28]^ the yield is severly compromised (6.3 ± 0.5 g when using a solvent mixture of 50% dwater/50% ethanol compared to 12.5 ± 0.2 g using 100% dwater for solubilization, and 100% ethanol absolute for precipitation). It might be due to the higher evaporation of ethanol when maintained at 45 °C for 1 h, instead of acting as a precipitation agent when maintained in solution at 25 °C. Furthermore, the waste increased when using the solvent mixture. The amount of 9.5 ± 1.2 g was produced compared 7.8 ± 0.1 g when using two isolated fractions of each solvent (**Supplementary Table 2**).

After ethanol, Pluronic F-127 (0.25% w/v) was added to the mixture as previously reported.^[28]^ Pluronic F-127 is a surfactant that has the role of stabilizing the gelatin core through hydrophobic interaction, aiding microparticle formation.^[36]^ At this stage, pH was often assessed, being at 6.4 ± 0.1. Even though previous work pointed pH of 6.25 to start precipitation,^[28]^ we observed that the solution was clear on this pH value, indicating absence of precipitation. Thus, our coacervation procedure was initiated by both altering pH and temperature of the system. We identified that lowering both pH to 5.8 and temperature to 25 °C, while keep stirring, resulted in an emulsion (**Figure 2D**), called here the micro-coacervate. At this stage, particles of gelatin and gum arabic were obtained and classified as microparticles^[29]^ with a mean diameter of 41 ± 8 µm (**Figure 1E**). Reducing the pH from 5.8 to 4.2, by slowly adding 1 M HCl, resulted in decreasing turbidity. At pH 4.2, the solution became clear again, demonstrating an absence of precipitation, and instead of microparticles, a hydrogel was formed overnight at 25 °C and 4 °C (**Supplementary** Figure 3). Interestingly, the behavior of the system is reversible when ranging from pH 4 to 6. This can be easily assessed by changing the turbidity of the solution from a transparent to an opaque one (**Supplementary** Figure 4), making easier any adjustment in pH. Together, these results indicate the following sequential conditions for enhanced micro-coacervation: 4:1 gelatin:gum arabic ratio dissolved in distilled water at 45 °C, addition of ethanol and pluronic, and adjustment of pH to 5.8.

Once identified above the best condition for enhance precipitation and yield of coacervates, the second step (macro-coacervation) starts when the emulsion is kept in static conditions at pH 5.8, 25 °C for 6 h. In previously reported work, the coacevation step was concluded at this stage, maintaining the coacervates at room temperature.^[28,37]^ We identified that a second temperature drop to 4 °C and overnight incubation resulted in exceptionally 7 times higher yields (**Figure 2D**) of embedding support (12.5 ± 0.2 g) when compared to maintaining it at room temperature (1.8 ± 0.3 g), per 100 mL of solution (**Supplementary Table 3**). The second temperature drop to 4 °C was also analyzed in terms of compactness of the coacervate and a much more compact coacervate was obtained after at 12 h on this temperature when compared to room temperature (**Supplementary** Figure 5). A quantitative characterization of the coacervate in all these steps is provided by measurements of turbidity as a function of pH (**Figure 2E**). At pH 6.5, the transmittance of the light is close to 100%, typical of a transparent solution. At pH 5.8, the transmittance of light is below 5%, due to the high turbidity of the emulsion and opaque charater of the microparticles. We observed that the macro-coacervate can be maintained hydrated on its fluid phase at 4 °C for at least two months without any negative effects. Once the fluid phase is removed, followed by a series of washing steps/centrifugation, the coacervates are ready for use at room temperature (**Figure 2F**). In contrast with previous work,^[37]^ which performs three sequential steps of washing with water, we do not recommend washing these support baths with water because the majority of protein-based bioinks are water-based. (**Supplementary** Figure 6). Thus, due to the high hydrophilic nature of water-washed support baths, the bioinks would diffuse and spread, imparing bioink printability. Thus, we washed the support baths with 20 mM CaCl_2_ due to its stabilizing effect resulting from additional linkages between gelatin chains that are often mediated by the Ca^2+^ ions.^[38]^ As less hydrophilic the bath is, the more it helps on holding water-soluble bioinks, such as PLMA and BSAMA, during printing. Thus we used a CaCl_2_-washed bath for our blood-based bioinks.

### Rheological characterization of supporting baths

The appropriate supporting bath for bioprinting must have a yield-stress and self-healing behavior, as well as a solid-like behavior at low shear stresses. These criteria are crucial because when the needle immerses during embedding bioprinting, it must move freely without any resistance from the support bath. If the bath offers resistance, such as when printing inside a full block made only of gelatin, the printed design is severely compromised **(Supplementary** Figure 7). Also, if the needle moves freely and stress is applied at higher levels than the yield stress, the supporting bath needs to accommodate the needle. Once the bioink is extruded and the needle moves, the supporting bath should recover its solid-like behavior and quickly close the groove created by the needle (**Figure 2J**). At the same time, the printed filament must be maintained in place with precise size fidelity and shape retention. In case the supporting bath does not recover immediately, the liquid bioink diffuses into the groove and is crosslinked in a totally spread and random conformation, not attending to the envisioned scaffold geometry. With these criteria in mind, the supporting baths produced by coacervation with defined gel strength of 150 and 250 gelatins with gum arabic were characterized by rheological assessment. According to amplitude sweep oscillatory assays, both baths present a solid-like behavior at room temperature (**Figure 2G**). For 250 bloom, the linear viscoelastic region (LVR) spans up to 4% strain, while for bloom 150, LVR was up to 6% strain. The supporting baths with 250 bloom gelatin were much stiffer than those with 150 bloom gelatin (G’= 889 Pa vs. G’= 156 Pa, respectively). This stiffness value for 250 bloom gelatin was similar to that reported in previous work^[28]^ (G’≈1100 Pa). In comparison, baths from the blend 50-120 gelatin eventually became liquid-like (G’’>G’) (**Supplementary** Figure 8). Frequency sweeps (**Figure 2H**) revealed similar results, with baths produced with 250 bloom being stiffer than those produced with 150 bloom. Analyzing flow sweeps (**Figure 2I**), supporting baths produced with 250 bloom were more viscous than those produced with 150 bloom (17.8 Pa·s vs. 2.5 Pa·s viscosity, respectively, for a 1 s^-1^ shear rate). These results suggest that gel strength interferes with microparticle formation, either in particle size, size distribution, or single-particle stiffness, as well as in their compaction state to form the bath.

To evaluate the self-healing properties, a three-step flow assay was performed (**Figure 2K**). Although both supporting baths recover the initial viscosity in <2 s when transitioning from the second-step high shear stress, to the third-step low shear stress, the 250 bloom gelatin bath recovered slightly faster. By comparison, the supporting bath with blend 50-120 gelatin took up to 9 s to recover the initial viscosity after being subjected to high shear stress (**Figure 2L**). Such results indicate that baths from blend 50-120 gelatin would not be suitable to provide support to the extruded ink. Hence, even though precipitation visually happens similarly with 150, 200, and 50-120 blend gel strength (**Supplementary** Figure 9), the self-healing properties are considerably different. Based on these findings, we conclude that gelatins with defined gel strength of 250 are the most suitable for preparing reproducible coacervates to be used as bioprinting support baths. Thus, the 250-bloom gelatin was used for all bioprinting studies and further characterization. These results also argue that the main properties of the chosen gelatin should be characterized and known before starting the coacervate procedure. We regularly observe that the characterization of gelatin is overall underestimated. Gelatin can vary considerably, being popularly called a black box material, but as presented, it can be easily characterized.

Interestingly, a recent report showed the versatility of using complex coacervates to obtain bioinks for embedding bioprinting, instead of the support bath as we envisioned.^[39]^ The authors exploited the properties of coacervate inks made of hyaluronic acid and chitosan and found parameters that can be used to fine-tune these inks for making them printable both in air and in water.^[39]^

### Formulation of BSAMA and PLMA hydrogels and crosslinking strategy

Once we developed the efficient support bath for low viscosity bioinks, we validated the bath for bioprinting blood derivatives as a biologically relevant example of low viscous liquid gels. The proteins PL and BSA were functionalized by the addition of methacryloyl groups to obtain photopolimerizable hydrogels, as previously described.^[8]^ Once lyophilized, PLMA and BSAMA were stable up to six months (**Figure 3A**). Preliminary screening of PLMA and BSAMA hydrogels was performed with the vial inversion test. Both 10% (*w/v*) in PBS solutions flowed immediately after vial inversion due to their high fluidity (**Figure 3A**). PLMA and BSAMA concentrations were increased to 30% (*w/v*) in PBS and the same instantaneous flow behavior was observed. Since increasing concentration did not make any of the solutions resist flowing, a fixed concentration of 10% (*w/v*) of PLMA and BSAMA was selected for the whole study. Ruthenium was used for photocrosslinking due to its cell cytocompatibility and water solubility.^[40]^ Once crosslinked, different PLMA and BSAMA hydrogels were obtained by varying intensity and exposition time (**Figure 3B**). Using visible light (455 nm) with intensities of 6, 11, and 21 mW/cm^2^, a minimum time of 90, 60, and 60 sec, respectively, was necessary to obtain hydrogels. However, hydrogels obtained using 6 and 11 mW/cm^2^ were extremely fragile and those obtained using 21 mW/cm^2^ were highly heterogeneous, presenting areas where the hydrogels were crosslinked and others were still liquid, thus clotting the syringe and disturbing the bioprinting process. When increasing the intensity of light to 46 mW/cm^2^ for 60 sec (**Figure 3C**), stiff and homogeneous hydrogels were obtained. These light parameters were used for subsequent experiments.

**Figure 3:**
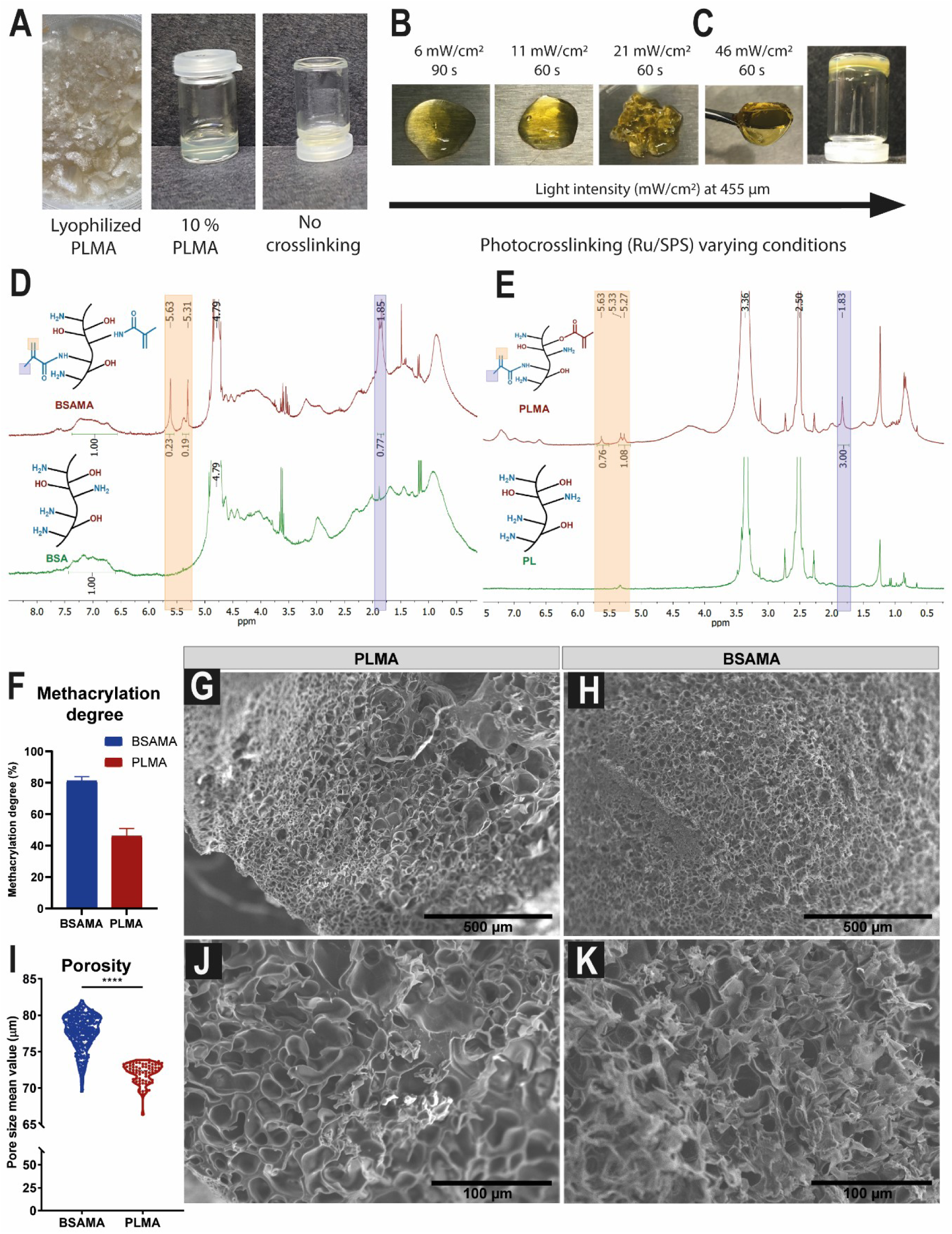
Properties and characteristics of PLMA and BSAMA photocrosslinkable hydrogels. **A)** Visual aspect of lyophilized PLMA samples and their high fluidity when solubilized in PBS before crosslinking. **B)** Hydrogels less or more fluid can be obtained by varying intensity and times of light exposition using the Ru/SPS photoinitiator system. **C)** Homogeneous hydrogel can be obtained using visible light at 455 nm, 46 mW/cm^2^ for 60 sec. **D, E)** ^1^H NMR spectra with distinctive peak characteristics of methacrylate groups for **(D)** BSAMA and **(E)** PLMA. **F)** Methacrylation degree of BSAMA and PLMA. Scanning electronic microscopy (SEM) micrographs of **G)** PLMA and **H)** BSAMA surface structures. **I)** Porosity of PLMA and BSAMA and respectives close-ups **J)** and **K**) evidencing details of the surface topography.

### Structural characterization of BSAMA and PLMA hydrogels

PLMA and BSAMA solutions as well as PL and BSA were characterized using ^1^H NMR; photoactive methacrylic groups were confirmed in both solutions. By analyzing BSA and BSAMA spectra (**Figure 3D**), it was possible to verify that BSAMA spectra contained specific peaks at around 5.3 and 5.6 ppm, which corresponded to the acrylic protons, and at approximately 1.8 ppm, corresponding to the methyl protons. Similarly, PL and PLMA were compared (**Figure 3E**), and the PLMA spectra exhibited the methacrylation characteristic peaks at 5.3 and 5.6 ppm of the acrylic protons and 1.8 ppm of the methyl protons.^[8]^ However, the methacrylation degree differed: 81.2 ± 2.7% for BSAMA, compared with 46.0 ± 4.9% for PLMA (**Figure 3F**). The significant difference in methacrylation degree is likely attributed to albumin being the main protein undergoing modification in PLMA. Albumin represents approximately 41% of the total protein content in PL.^[8]^ We hypothesized that BSAMA will lead to more stable printed filaments along with higher printing fidelity compared to PLMA, thus it was crucial to characterize these solutions as bioinks and respectives hydrogels.

Scanning Electron Microscopy (SEM) analysis shows the surface microporosity of PLMA (**Figure 3G**) and BSAMA (**Figure 3H**) freeze-dried hydrogels. **Figure 3I** shows that both protein-based hydrogels present a random distribution of pores where the PLMA hydrogel (**Figure 3J**) present less regular ones in terms of structure and size distribution than BSAMA hydrogel (**Figure 3K**).

### Rheological characterization of PLMA and BSMA solutions and hydrogels

BSAMA and PLMA precursor solutions were both characterized by rheological measurements. The shear rate sweeps (**Figure 4A**) demonstrated that both solutions exhibit shear-thinning behavior. As expected from the vial inversion test, their viscosity was extremely low. Suitable inks for extrusion-based printing usually present viscosity values between 3×10^-2^ and 6×10^4^ Pa·s,^[41]^ but BSAMA and PLMA have shown around < 10^-3^ Pa·s in viscosity, which means that both solutions present close viscosity values to water (10^-3^ Pa·s, as a Newtonian fluid)^[42]^. BSAMA and PLMA precursor solutions were submitted to amplitude sweeps to determine their linear viscoelastic region (LVR). Both solutions presented again a very soft nature, with a LVR in the 3–20% strain range (**Supplementary** Figure 10).

**Figure 4:**
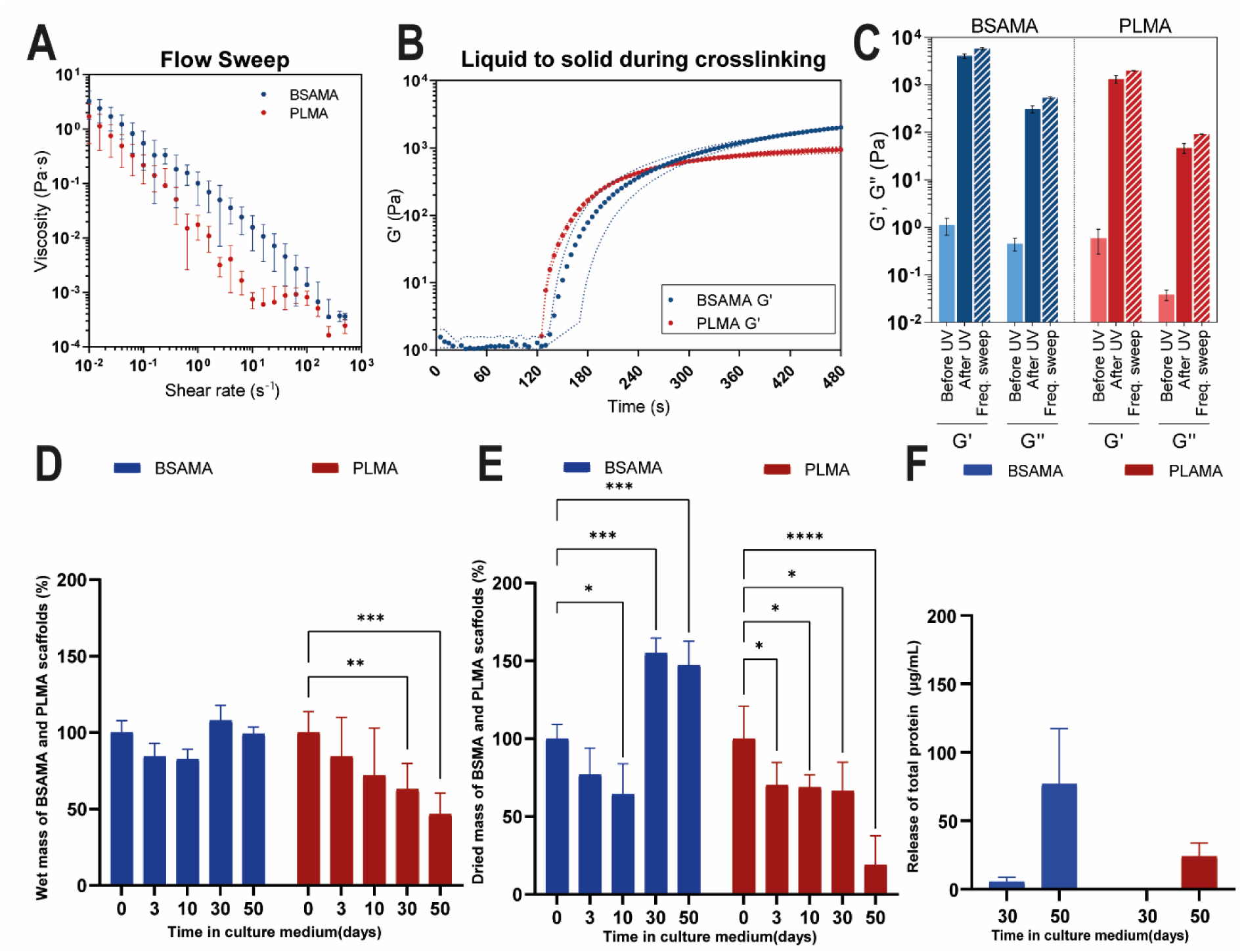
**A)** Flow behavior of BSAMA and PLMA bioinks. **B)** Photocrosslinking kinetics of BSAMA and PLMA hydrogels. **C)** Storage modulus (G‘) and loss modulus (G‘‘) of BSAMA and PLMA before and after crosslinking, and as calculated from frequency sweep oscillatory assays. **D)** Swelling of BSAMA and PLMA hydrogels as measured by wet mass compared to day 0 over 50 days in culture medium. **E)** Mass change of BSAMA and PLAMA hydrogels over 50 days in culture medium, as measured by dry mass relative to day 0. **F)** Release of BSAMA and PLAMA components in the surrounding microenvironment at day 30 and day 50 in culture.

Next, we characterized crosslinking kinetics by photorheology. BSAMA and PLMA solutions were submitted to an oscillation assay, and their G’ and G’’ were monitored throughout time, before and after irradiation (***Figure 4B***). After irradiation, their storage moduli increased logarithmically and eventually stabilized, indicating crosslinking. The stabilization time was estimated to be 63 ± 3 sec for PLMA hydrogels, and 213 ± 16 sec for BSAMA hydrogels. It is worth emphasizing that such crosslinking times are long in the context of bioprinting. Hence, printing them through conventional “on-air” standard extrusion was unattainable, reinforcing the need for a support bath to allow shape retention. Interestingly, while BSAMA hydrogels crosslinked slower than PLMA, they were stiffer when the crosslinking process was complete: BSAMA hydrogels demonstrated a G’= 4.08 ± 0.03 kPa and G’’= 0.31 ± 0.02 kPa, while PLMA hydrogels presented G’= 1.33 ± 0.01 kPa and G’’= 0.05 ± 0.0001 kPa (**Figure 4C**). Such moduli are close to those calculated through frequency sweeps, with BSAMA hydrogels displaying G’= 5.78 ± 0.29 kPa, G’’= 0.53 ± 0.03 kPa compared to PLMA hydrogels of G’= 1.99 ± 0.002 kPa, G’’= 0.091 ± 0.001 kPa (**Supplementary** Figure 11). This difference can be correlated to their composition since BSAMA has a higher methacrylation degree than PLMA (**Figure 2F**), hence a higher crosslinking density. Additionally, BSA is a large molecular weight protein, in comparison with the remaining components of PL based on growth factors, cytokines, peptides, and lipid-binding molecules, among others.

### Swelling index, mass change, stability, and release of PLMA and BSAMA hydrogels over 50 days

Finally, to verify if BSAMA and PLMA bioinks were capable of supporting long cell culture experiments, the swelling effect, mass change, release of components and consequently stability were assessed using crosslinked droplets of these materials (10% *w/v* in PBS) maintained in medium culture over 50 days. Surprisingly, for the BSAMA ink there was neither swelling effect nor hydrogel degradation since the wet mass was maintained over time (***Figure 4D***). BSAMA proved to be a very stable photocrosslinkable system for 50 days (100 ± 8 % at day 1 and 99.2 ± 4 % at day 50, **Supplementary** Figure 12), thus holding great potential as a drug delivery system (DDS).^[43]^ Several albumin-based DDS have already received regulatory approval, suggesting that this system will continue to be a useful method for future endeavors.^[43]^ Regarding PLMA, there is also no swelling effect, but the mass decresead over time, signifying degradation (***Figure 4D***). The degradation of PLMA hydrogel started only at day 30, indicating it is a stable choice for stem cell differentiation procedures that generally span 28 days.

As another method to assess mass change, we measured the dried mass (**Figure 4E**). Similar to the wet mass, BSAMA showed a loss until day 10, after which the mass significantly increased until day 50. The mass loss and gain observed with BSAMA may be due to increased retention of culture medium components, similar to that observed when using polysaccharides.^[44]^ BSA is known as an efficient blocking agent since it binds to at least 40% of the circulating calcium and acts as a carrier of several molecules, such as long-chain fatty acids, ions, hormones, and FBS components.

In contrast, PLMA hydrogels showed a rapid decrease to 70.2 ± 15% at an early time (day 3), which was maintained around this level (66.4 ± 19%) until day 30, emphasizing its stability over 30 days. At day 50, less than 20% of the dried mass of the platelet lysates could be found, indicating degradation. Such degradation can be advantageous for a cellular microenvironment where cell-recruiting and pro-regenerative molecules are released as the platelet lysates degrade. In fact, the release of proteins was assessed (*Figure 4F*), with 77% and 24% of BSMA and PLMA components, respectively, identified in the supernatant at day 50.

### Reproducibility and cytotoxicity tests of PLMA hydrogels using hMSCs

Aiming to assess PLMA reproducibility for cell culture using human mesenchymal stem cells (hMSCs), PLMA batches of three independent productions were evaluated. There was no difference in the metabolic activity of hMSCs after 24 hours of culture in these three PLMA batches (**Supplementary** Figure 13). This is an important finding, especially considering highly variable human blood components, thus emphasizing the high control of extraction and production of these materials. Considering that some free methacrylate groups could be in excess in PLMA due to the functionalization procedure, we also analyzed the metabolic activity of hMSCs encapsulated in PLMA/Ru solutions in the three PLMA batches. There was no significant difference in the metabolic activity of PLMA hydrogels when crosslinked 1 or 45 minutes after preparation (**Supplementary** Figure 14). These findings emphasize that the PLMA system is not cytotoxic and presents a safe choice regarding hMSC cell culture, corroborating with previous reports.^[45,46]^

### Bioprinting PLMA and BSAMA hydrogels using supporting baths

Next, we assessed the printability of the PLMA and BSAMA hydrogels in the gelatin-gum arabic supporting bath developed by CLADDING. Using the same printing parameters, it was evident that well-defined filaments could be obtained using the supporting bath comprising 250 bloom gelatin (**Figure 5A**), while no defined structure close to a filament could be obtained when using standard bioprinting techniques (**Figure 5B**). From a processing point of view, **Figure 5A** clearly shows the high reproducibility of our developed supporting bath and the small amount that is required for bioprinting each scaffold. Since the bath doesn’t flow on the plate, only the necessary amount to cover the envisioned design is needed, an advantage compared with other support baths that require covering up the whole well. All these aspects offer a significant improvement for scaling-up scaffold production when using embedding bioprinting.

**Figure 5:**
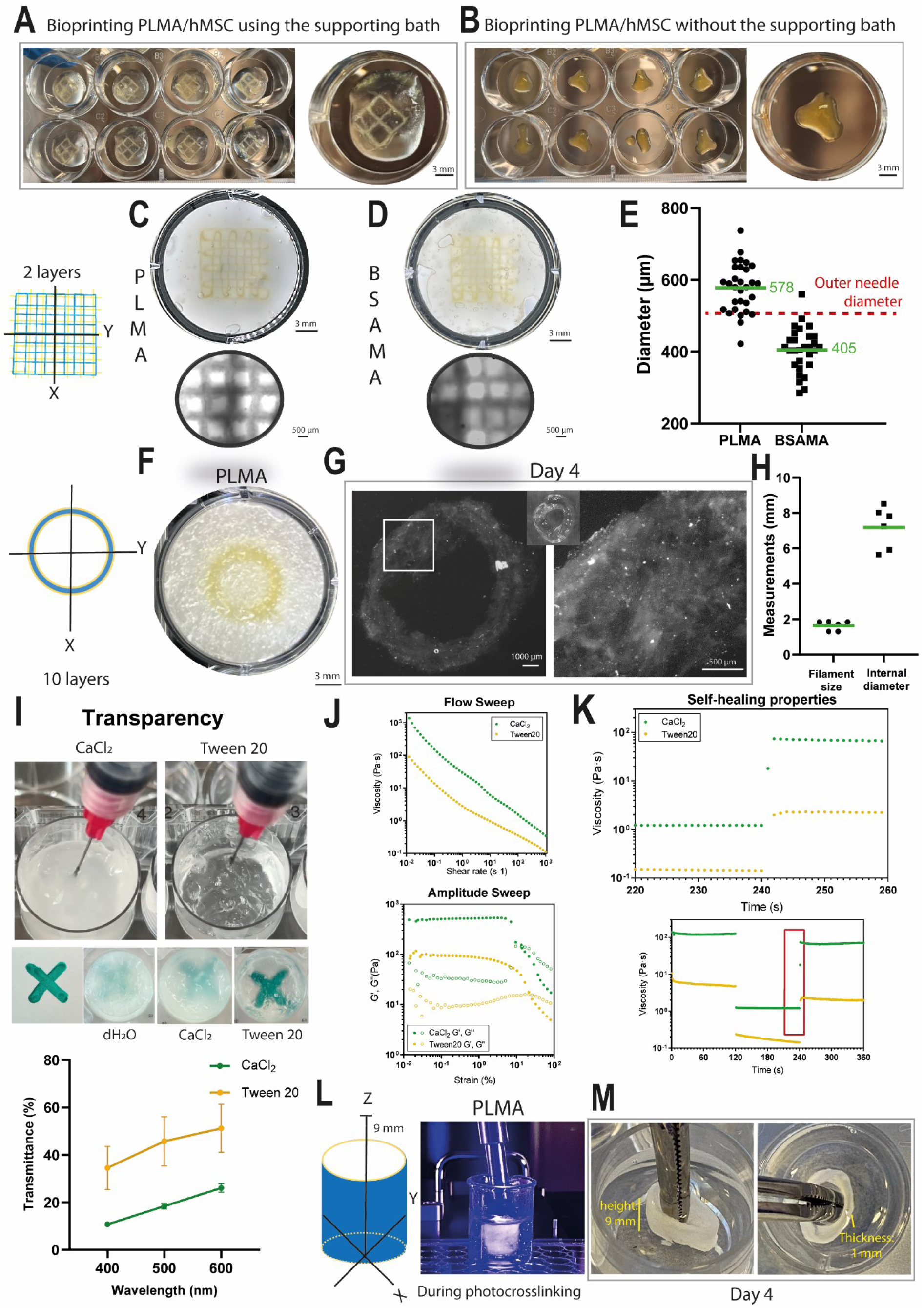
Bioprinting PLMA/hMSCs and BSAMA/hMSCs bioinks in the gelatin-gum arabic supporting bath developed by CLADDING. The same bioprinting parameters were applied when bioprinting PMLA/hMSCs. **A)** inside the 250 bloom gelatin supporting bath and **B)** in the air without the supporting bath. Well-defined filaments with ideal layer-layer attachment were obtained when bioprinting **C)** PLMA/hMSCs and **D)** BSAMA/hMSCs. **E)** Comparison of the obtained filament size when bioprinting PLMA/hMSCs and BSAMA/hMSCs bioinks using a needle of 520 µm outer diameter. **F)** Bioprinted rings composed of 10 layers of PLMA/hMSCs. **G)** PLMA/hMSCs rings were stable in culture media for up to 4 days with no delamination or degradation. **H)** Filament size and internal diameter of 10-layer PLMA/hMSCs rings. **I)** Washing with 5% v/v Tween 20 resulted in transparent support baths. In comparison, washes with Tween 20 and CaCl_2_ (bottom row, right images) improved transparency when compared to distilled water (bottow row, left image). Quantitatively, the double of the amount of light was transmitted in support baths washed with Tween 20 when compared with CaCl_2_. **J)** Flow behavior of washed supporting baths using Tween 20 and CaCl_2_. **K)** Self-healing behavior of washed supporting baths using Tween 20 and CaCl_2_.**L)** Bioprinting of 10 layers moving in Z for obtaining cylinders. **M)** Exhibition of a stable cylinder with 9 mm of height and 1 mm of thickness after the support bath removal.

In terms of printability, for embedding bioprinting the envisioned filament size is defined based on the outer diameter of the needle, since this is the size of the groove created in the supporting bath when the needle moves. Using a needle of 520 µm outer diameter (25 gauge), we obtained well-defined filaments of 578 ± 68 and 405 ± 60 µm with PLMA (**Figure 5C**) and BSAMA (**Figure 5D**), respectively. The layers attached very well to each other proving tha the supporting bath does not impair the attachment in between filaments. As a matter of comparison, the same experiment was performed in a more commonly used commercial support bath (LifeSupport, FluidForm^TM^, **Supplementary** Figure 15). Using the same 25 gauge needle, there was no significant difference in terms of filament size. However, a clear advantage of our approach as that it reproduced much better the intended design in terms of printing fidelity, due to lower bioink spreading and improved adherence between layers. Pore printability index, pore area and pore lateral size were calculated to characterize the difference in inner geometry of the constructs (**Supplemetary Figure 15**). CLADDING bioprinting showed a better performance and much lower variability than the commercial support bath. As previously demonstrated when comparing oil and water-based baths, shape definition is achieved when the supporting bath minimizes the electrostatic interaction at the bioink-bath interface during printing. ^[47]^ These findings show that our self-healing support bath effectively minimized these electrostatic interaction since very accurated filaments that nicely attach to each other were obtained with high fidelity to the designed conditions (**Figure 5C-E**). Challenging the system with a much smaller diameter as technically possible with current available needles for embedding bioprinting (310 µm outer diameter, 30 gauge/150 µm of internal diameter), filaments of 142 ± 36 µm with PLMA were obtained in an intact structure **(Suplementary Figure 16**).

Also, correlating with the methacrylation degree of BSAMA and PLAMA and the viscosity of the inks previously discussed (**Figure 3F**), it is evident that the higher viscosity and the methacrylation degree of BSAMA led to higher compaction of the hydrogel network and smaller diameter of the printed filaments.

Next, we progressed to bioprinting circular models using PLMA, such as rings and cylinders since they lack structural pillars that help during standard bioprinting, and due to the existence of many structures like these in the human body, such as vascular and neural tissues.^[48,49]^ Stable rings fully composed of PLMA were obtained when bioprinting multiple passes of 10 layers to generate thicker structures along with simultaneously photocrosslinking (**Figure 5F and Supplementary Video 1**). When we challenged the system by switching bioprinting multiple passes for bioprinting 10 circumferences in a continuous spiral with a pitch equal to the fiber diameter, the filaments coalesced since the filament size of the whole rings was 1.6 ± 0.3 mm, instead of 578 ± 68 µm of a single layer. Coalescence is a common phenomenon when bioprinting multilayered structures, as already reported by our group.^[44]^ We believe that this result is still suitable for the high accuracy of the system since bioprinting 10 layers in a continuous spiral requires a longer printing time, thus more spreading is also expected. These 10-layer rings of 7 mm in core diameter and 9 mm of height were bioprinted and maintained up to 4 days (**Figure 5H**). After 4 days in culture media, the PLMA rings were still stable and delamination between layers was not observed, proving that the supporting bath did not impair the attachment between layers also in circular models (**Figure 5G**). As a proof of concept for more stable yet low viscous liquid hydrogels such as PEG-based bioinks (**Supplementary** Figure 17), rings of 4 mm of core diameter could be successfully obtained and maintained for up to 20 days.

Overall, we could draw a correlation between methacrylation degree, rheological properties and performance during bioprinting when using BSAMA and PLMA bioinks. BSAMA bioinks showed viscosity values 2× higher (for shear rates relevant in extrusion printing) and methacrylation degree 1.8× higher than PLMA. Due to its higher viscosity and crosslinking density, BSAMA showed superior printing fidelity, i.e. smaller filaments in size. Nevertheless, photocrosslinking of BSAMA was slower than PLMA, most probably because of the higher density of methacrylate groups in BSAMA and also due to its globular nature, which spatially impairs in a quick crosslinking, but increases both network density and consequently elastic modulus. PLMA bioinks showed a faster photocrosslinking, since it has less than half of photocurable moieties available when compared to BSAMA, as well as due to steric hinderances of other proteins in PLMA. This correlation should be taken into account for the envisioned mechanical properties of PLMA- and BSAMA-bioprinted constructs. We believe that both blood components hold great potential to be applied to a range of tissue engineering applications, and the choice should consider mainly the different levels of stiffness, stability, and biological relevance.

### Washing supporting bath as a function of transparency and hydrophilicity

Having observed that constructs were stable for up to 4 days, we intended to prolong this time to ensure a long-term cell culture process. Thus, we focused on improving the photocrosslinking step. Analyzing the absorbance of the blue light (455 nm) in the support bath in different depths, we observed that the height of the bath support was directly correlated to the absorbance of 455 nm light (**Supplementary** Figure 18). Indeed, this observation is consistent with a common phenomenon of light interaction with particles, mostly related to absorption and scattering, being the limiting height determined using the Lambert-Beer law.^[50,51]^ Interestingly, this same relationship was previously reported when irradiating blue light (460 nm) in media culture, showing that this phenomenon occurs when light is irradiated in different solutions than air.^[50]^ Thus, aiming to overcome this absorbance and improve our support bath, we investigated compounds that could reduce the turbidity of the bath by promoting self-organization on the gelatin–gum arabic chains. We selected NaSO_4_ and NaPO_4,_ based on the Hofmeister series and strategies for the salting-out effect,^[52]^ aiming to enhance the hydrophobic effect. We also selected Tween 20, as a surfactant with a hydrophilic head followed by a hydrophobic alkyl tail and often used to promote self-organized adsorbate layers, since it might have the ability to increase the hydrophobicity of the system.^[53]^ Also, Tween 20 has a particular resistance against protein adsorption, thus it has been widely used as a protein-resistant coating, as well as to also increase stability.^[53]^

Using NaSO_4_ and NaPO_4_, no improvements could be seen and the opacity remained the same. Comparing the CaCl_2_-washed based as abovementioned and now Tween 20, we observed a biphasic system right after the three washing steps with both washing solutions. The solvent phase was transparent, demonstrating that it was efficiently isolated from the coacervate phase (**Supplementary** Figure 19). Using CaCl_2_, a less opaque coacervate could be generated compared to using only water for washes (**Figure 5I**). When performing a proper wash using Tween 20 (5% v/v), the support bath became transparent (**Figure 5I**). Quantitatively, the double of the amount of light were transmitted in support baths washed with Tween 20 when compared with CaCl_2_ (**Figure 5I**). This is a great result for enabling embedding bioprinting using photocrosslinkable materials. We hypothesize that Tween 20 stimulated self-organization of the polymer chains of gelatin and gum arabic, yielding a transparent bath. In corroboration to the herein presented findings, previous work that studied the lauric arginate ester-carrageenan coacervate for food grade applications showed that once SDS or Tween 20 were mixed into the coacervates, a reduction in turbidity was noted. The authors hyphotesized that turbidity decreases in presence of these agents due to a reduction in the overall charge of the system, weakening the polymer-protein complexation and redispersing them into the solvent.^[54]^ Since Tween 20 is a detergent compound, we immediately assessed hMSC viability to identify if this washing step is a safe option. No cytotoxicity effect on hMSCs was observed (**Supplementary** Figure 20).

Finally, we investigated if these washing steps using CaCl_2_ and Tween 20 could modify the rheological properties of the supporting bath. Thus, coacervate phases were collected and washed three times with 20 mM CaCl_2_ (as initially optimized) or 5% v/v Tween 20 (as now proposed for transparency). Analyzing flow sweeps (**Figure 5J**), it can be seen that baths washed with Tween 20 presented a lower viscosity than those washed with CaCl_2_. Likewise, from amplitude sweep oscillatory assays, baths in Tween 20 were softer than those in CaCl_2_ (**Figure 5J**), but both presented a solid-like behavior. Regarding their self-healing behavior, baths washed with Tween 20 recovered almost completely from high shear stress in < 1 s (despite stabilizing only after 3 s), having just a slightly faster recovery than baths washed with CaCl_2_ (**Figure 5K**). **Supplementary Video 2** shows the self-healing behaviour while the baths were submitted to intense shear stresses by the noozle. However, it is important to note that for Tween 20 the bath’s viscosity decreased slightly when submitted to high shear rates (100 s^-1^) for the 2 min tested, unlike for CaCl_2_. This decreased viscosity may indicate disruption of the microparticle structure when under high shear stresses throughout time.

Finally, after washing our supporting baths with Tween 20 as an efficient strategy to overcome the limited penetration depth of the 455 nm wavelength, we challenged the system by printing larger structures, such as cylinders, by printing 10 layers of PLMA bioink moving in Z direction(**Figure 5L**). Stable and taller cylinders of 9 mm in height and 1 mm in thickness were obtained (**Supplementary Video 3).** The big structures also retained their intact shape after support bath removal (**Figure 5M**). The filament size was 50% smaller compared to that when maintaining the Z-axis fixed in the same position (**Figure 5G**). This smaller size is probably due to the lower coalescence effect when the Z-axis moves up continuously, as a result of printing the bioink in a suspended free-form approach.

### Analyzing cellular behavior of hMSCs after encapsulation and bioprinting with PLMA bioink

Finally, we investigated the behavior of hMSCs when encapsulated in PLMA hydrogels, right after bioprinting grid constructs and up to 7 days in culture, especially in terms of cell viability, metabolic activity, cell morphology, cell spreading, and attachment, which altogether can offer an understanding of the cell-recruiting properties of PLMA bioink. The cell density employed here for hMSCs in single culture (4 million hMSCs per mL of PLMA) proved to be satisfactory regarding metabolic activity and cell distribution. Right after bioprinting (**Figure 6A**), a highly homogenous distribution of cells can be observed throughout the filament. Cells were in a spherical form and well dispersed in single cells. On day 1 after bioprinting (**Figure 6B**), cells spontaneously started to aggregate and form spheroids. This could be due to the presence of the melted support bath within the constructs in which the chains of gum arabic impaired cell attachment on the PL components, or most probably due to the lack of enough bonding sites for cell attachment since it is a 10 % PLMA solution. Due to media refreshment, the supporting bath was usually depleted around day 4 (data not shown). On day 4 after bioprinting (**Figure 6C**), a high increase in cell population could be observed, and the cells started to migrate within the PLMA network and aggregated into spheroids smaller than 200 µm. On day 5 after bioprinting (**Figure 6D**), many spheroids began to disassemble into single cells and the remaining spheroids were of variable sizes, ranging from tiny (43.6 ± 8 µm), small (101.2 ± 19 µm), medium (264.9 ± 33 µm), big (365.8 ± 51 µm) and large (516.5 ± 49 µm). At day 7, cells showed high viability (**Figure 6E–F**) either attached in the PLMA filament (**Figure 6E**) or in aggregates (**Figure 6F**). Metabolic activity also demonstrated a significant enhancement of 1.5× at day 4 compared to day 1 (**Figure 6G**). Thus, we analyzed if bioprinting PLMA using culture medium as a solvent instead of PBS would result in increased metabolic activity from day 1 to 7 (**Figure 6H**). However, no statistical significance was observed when comparing the culture medium and PBS. Also, the stability of droplets obtained with PLMA/PBS or PLMA/culture medium was assessed, and with both solvents the droplets were stable for 40 days (**Supplementary** Figure 21). Thus, we conclude that both PBS or culture medium can be used for bioprinting PLMA.

**Figure 6:**
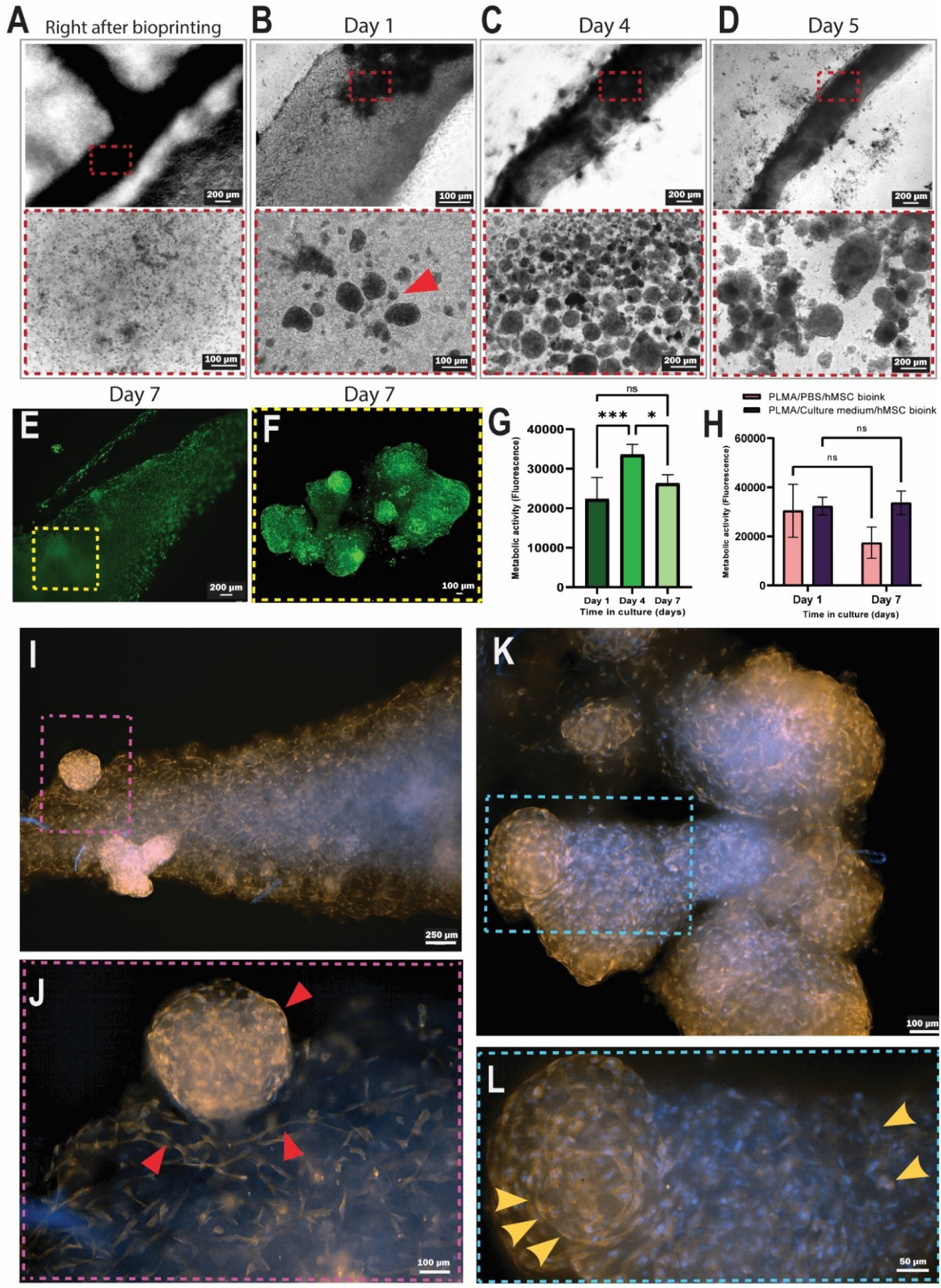
Cellular behavior in CLADDING 3D bioprinted constructs. **A)** Right after bioprinting, hMSCs showed a high homogenous distribution throughout the filament. **B)** On day 1 after bioprinting, cells spontaneously aggregated and formed spheroids (red arrow). **C)** At day 4, the cell population increased and spheroids could be seen. **D)** At day 5, spheroids dissembled into single cells. (**E, F**) Staining of live (calcein AM, green) and non-viable (EthD-1, red) cells at day 7 showed high cell viability in the **E)** PLMA filament and in **F**) spheroids. (**G, H)** Metabolic activity of hMSCs in bioprinted PLMA after 7 days, printed using PBS or culture medium (significant differences *** and * and non-significant, ns, were determined by Tukey’s multiple comparisons test). **I–L**) F-actin (orange) and cell nuclei (blue) staining demonstrated the ability of hMSCs to grow and spread both as **I)** single cells and as spheroids throughout PLMA filaments, **J)** in the spheroid-attached interface region (red arrows), and **K-L)** in fused spheroids. F-actin filaments formed circular patterns in the case of the spheroids (yellow arrows).

PLMA bioprinted constructs at day 7 were stained for cell nuclei and F-actin filaments, showing a highly dense cellular network. HMSC tended to align and spread on the borders of the PLMA filaments compared to the cores, where they spread less and were distributed in a more random pattern (**Figure 6I–L and Supplementary** Figure 22). Interestingly, hMSCs showed their robustness in either perfectly attaching and stretching on the PLMA filament, as well as in growing in 3D conformations when combined with PLMA hydrogel. The results are consistent with the previous observation of the good cell metabolic activity, proliferation and spreading ability of hMSCs when encapsulated in PLMA hydrogels, but not bioprinted.^[55,56]^ The interface region clearly showed the same morphological pattern between the attached and flat cells within the outer layer of the spheroid (red arrows, **Figure 6J**). Remarkably, the spheroids remaining at day 7 showed the capability to fuse with each other to form a tissue-like structure (**Figure 6K**). After fusion, an interesting ECM remodelling was observed (yellow arrows) as a result of the differences on F-actin distribution while in spheroids or in the forming tissue-like structures (**Figure 6L**).

Finally, using all the developments on the coacervation with gelatin-gum arabic presented herin, we aimed to validate if the self-healing and transparency capability of the CLADDING support bath would allow bioprinting low viscousity blood derivatives in mild conditions for bioprinting sensitive cell lines in a co-culture system, such as endothelial cells (hUVECs) with hMSCs.

Different structures containing a high cell concentration in co-culture (10 million/mL) of hUVECs (8 million/mL) and hMSCs (2 million/mL) were bioprinted while increasing complexity (**Figure 7A**). Circular filaments (without structural pillars), suspended tubular structures and porous constructs were obtained with high printing fidelity and could be easily maintained in cell culture for over a month (**Figure 7A-B**). After support bath removal, all constructs showed intact structural stability (**Figure 7C and Supplementary Video 4**). This is an excellent improvement of this technology since we started the process bioprinting only a liquid solution of platelet lysates highly concentrated with sensitive cells, without any polymer, thickener, molding or chemistry modification to increase its viscosity. At day 1 after bioprinting (**Figure 7D**), hUVECs and hMSCs were uniformally distributed throughout the PLMA filaments. Interestingly, they did not form any spheroids as observed using a single culture of hMSCs (**Figures 6A-D**). This might be probably due to the higher cell concentration used in the co-culture system compared to the single system (10 million *versus* 4 million) and due to the increased sites for cell bonding in the PLMA constructs since we used a higher PLMA concentration for the co-culture system (30% PLMA) compared to the hMSC culture only (10% PLMA). At day 7 (**Figure 7E**) cells showed a satisfactory spreading and a higher amount of hUVECs were noticed on day 7 compared to day 1. When bioprinting taller cylinders, the perfect layer-by-layer deposition could be clearly seen, with excellent attachment and no delamination while maintaining an appropriate porosity for oxygen and nutrient diffusion (yellow arrow). At day 14 (**Figure 7F**), cells still showed a homogenous distribution; the structure and porosity of the filaments were preserved. The cell viability of the whole period was satisfactory since high cell viability was observed at day 1 and 14 (**Figures 7G-H**), with a low amount of dead cells on the border of the filament at day 1 (**Figure 7G**), which is common due to the bioprinting process, and excellent cell distribution and spreading at day 14 (**Figure 7H**). Thus, using this enhanced support bath with self-healing and transparency capability, bioprinting low viscousity blood derivatives in mild conditions containing a high concentration of sensitive cells in co-culture proved to be feasible and reproducible.

**Figure 7:**
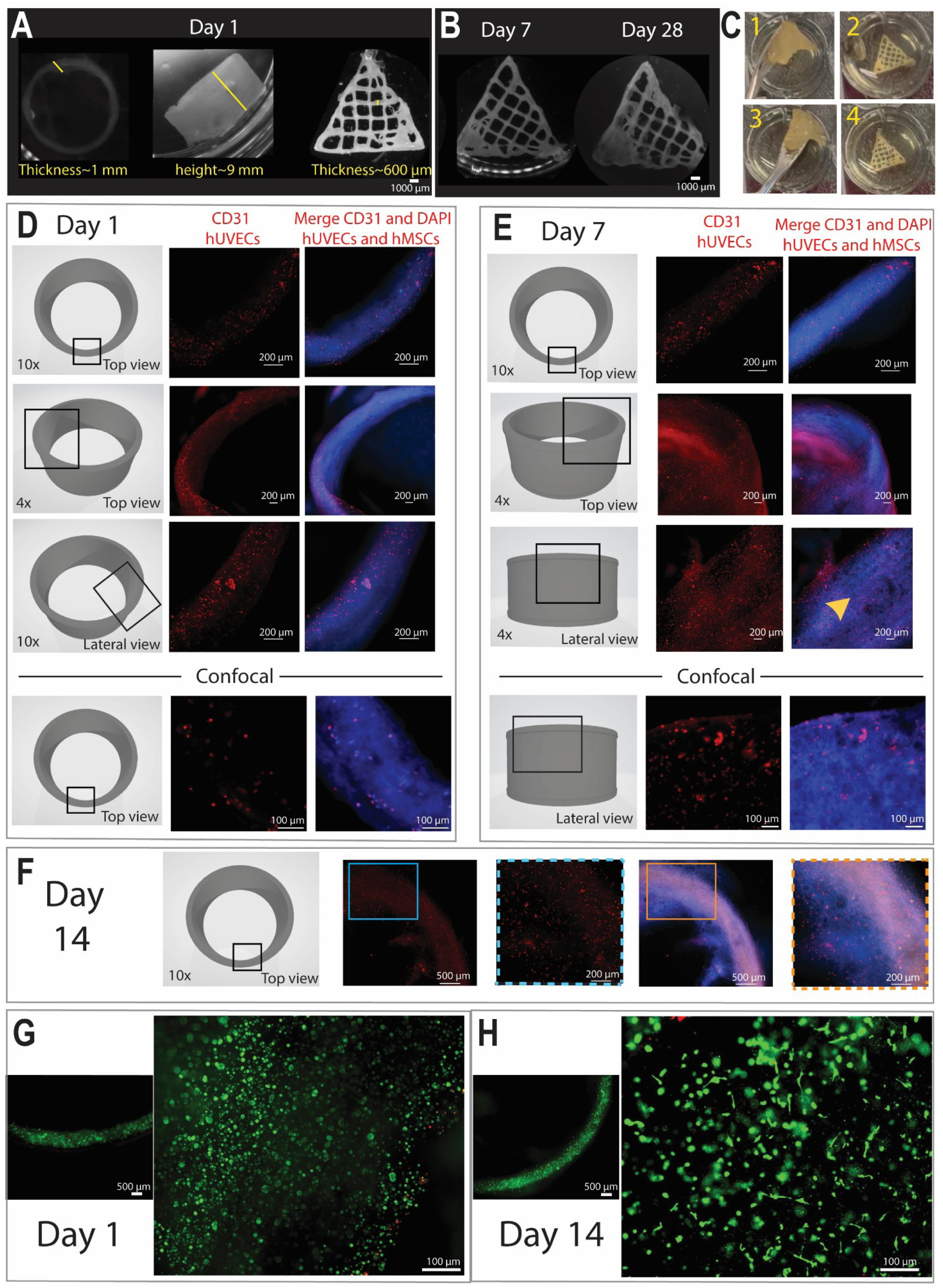
Bioprinting PLMA with co-culture of hMSC and hUVECS, in the gelatin-gum arabic supporting bath developed by the CLADDING method, while. **A)** increasing complexity of the structures: circular filament (left), suspended tubular structure (middle) and porous constructs (right). **B)** Structural stability of porous bioprinted constructs after removal of the support bath for 7 and 28 days. **C)** Repeated handling with spatula demonstrates high structural stability of porous bioprinted constructs after support bath removal. **D)** Day 1 after bioprinting PLMA with co-culture of hMSC and hUVECs in a high concentration of 10 million cells per mL of hydrogel. Staining of hUVECs using the endothelial cell-specific marker CD31 (red) and all cells (Dapi, blue) and the upcoming **E)** day 7 and **F)** day 14. **G, H)** Staining of live (calcein AM, green) and non-viable (EthD-1, red) cells at day 1 and 14 showing high cell viability in a PLMA filament. The cells showed a spherical morphology at day 1 and an efficient streatching and spreading at day 14.

Previous studies have described PL as a cocktail of human growth factors and bioactive molecules involved in cell growth, proliferation and recruitment.^[2,8]^ From proteomic analysis, vascular endothelial growth factor (VEGF), transforming growth factor (TGF-β), platelet-derived growth factor (PDGF), fibroblast growth factor (FGF), and epidermal growth factor (EGF) were ranked as high effectors on human PL activity, suggesting a major role for PLMA in healing damaged tissues/organs.^[2]^ A previous study identified that VEGF could be either released in a sustained manner or stay immobilized within a PLMA hydrogel for up to 10 days.^[8]^ VEGF is known for its critical role in cell recruitment of stem cells and immune cells to sites of neovascularization,^[2,12]^ thus we believe that it is one of the main growth factors responsible for aiding in the maintenance of hMSCs and hUVECs in the bioprinted constructs.^[2]^ Along with VEGF, fibrinogen, a key element in PRP and PLMA hydrogels, is also released in a sustained manner, but in very little amounts, suggesting that it plays a structural role in the PLMA hydrogels.^[8]^ The release of bioactive molecules from PLMA hydrogels permits the culture of cells in PLMA scaffolds in xeno-free conditions.^[46]^ Taking all into account, it is clear that bioprinted PLMA in a single culture of hMSC or co-culture of hMSCs with hUVECs showed a high-density cellular network with excellent cell adhesion, fast proliferation, and high cell viability. This demonstrates the bioactive properties of the PLMA-based bioink and emphasizes its potential for advancing tissue engineering and regenerative medicine applications.

## Conclusion

A universal approach to bioprint high low-viscosity bioinks through a multiscale coacervation procedure was herein presented, which we termd CLADDING bioprinting. The main physical principles behind coacervation of embedding bath and the parametric fabrication window to produce supporting baths by coacervates using gelatin-gum arabic was elucidated.

Gelatin properties were essential to be characterized, since the use of gel strength (bloom) as the individual selection criteria is not enough for coacervation procedures. The examination of molecular weight distribution, isoelectric point, ionic content, along with bloom altogether stands as an effective selection criteria. Several gelatin types were excluded for further testing because of the inappropriate performance on these parameters. An efficient coacervation procedure using different classes of gelatin (defined and non-defined) and gum arabic was developed to obtain microparticles, that showed a strong tendency to aggregate forming a typical colloid, ultimately composing a multiscale support bath for embedding bioprinting. The optimized support bath presented an ideal viscosity and excellent self-healing properties. A series of washing steps using calcium chloride and Tween 20 generated support baths with improved stability and transparency for photocrosslinking. Thus, the main novelty our study brings is the stablishment a method to produce a self-healing, transparent and multiscale supporting bath. This support bath is solving a key problem as it opens the spectrum of low viscosity photocrosslinkable materials that can now be bioprinted while encapsulating sensitive cells in single or co-culture systems such as endothelial and mesenchymal stem cells.

Methacrylated platelet lysates and serum albumin hydrogels were combined with hMSCs and hUVECs to obtain bioactive bioinks. Yet, both these bioinks prsented unsuitable rheological properties for standard bioprinting. BSAMA could be efficiently used as a standardized alternative to optimize processing involving photopolimerizable blood derivatives, whereas PLMA bioink as a cocktail of human growth factors and bioactive molecules involved in cell growth, proliferation, and recruitment. Throughout embedding bioprinting in our self-healing, transparent and multiscale supporting bath, bioprinted constructs fully composed of PLMA, hMSCs in single culture or co-culture with hUVECs were obtained, and showed excellent cell viability, increasing metabolic activity and high-dense cellular network. It is the first time that low viscousity blood derivatives are bioprinted in a co-culture system with sensitive cells without the use of any thickner or molding. After support removal, constructs showed excellent integrity for handling for a future implantation step and enough stability in culture media over a month, while maintained their bioprinted architecture.

Overall, our results showed a new improved method for embedding bioprinting of highly biocompatible xeno-free material with high fluidity without any additive to increase viscosity. This finding has great significance since it is based on human blood derivatives that can be applied to tissue engineering of many types of organs. With the increasing trend on the use of standardized clinical grade human components, this work represents an advantage over similar materials typically obtained from xenogenic sources, such as GelMA, ColMA, and ECM bioinks. Due to the universality of this approach, a high number of non-printable human proteins and peptides can be tested. The present work offers substantial progress for processing clinical grade human components.

## Material and Methods

### Physicochemical characterization of different gel strength gelatins regarding molecular weight distribution, isoelectric point, conductivity, and ion content

For the investigation of different types of gelatin, defined ones with gel strength of 50, 100, 150, 200, 250 (all called type I), from Rousselot were used. The non-defined one with gel strength of 50–120 (called type II) from Sigma-Aldrich (G6650) was used.

The molecular weight and distribution were measured using size exclusion chromatography (SEC) on a liquid chromatography (LC) machine, connected with a UV detector (Agilent HPLC, 1260 Infinity series). The 2% (w/v) gelatin solution of different types were further diluted in the mobile phase consisting of PBS saline buffer at pH 7.4 (Carl Roth, 1111.2) with added sodium dodecyl sulphate (SDS, Carl Roth, 0183.3). Further, a benzoic acid internal standard (Carl Roth, P738.1) was added to the sample. An aliquot of 20 µL of the sample was injected into the machine and carried at a flow of 0.5 mL/min by the above-mentioned mobile phase to a TSKgel PWXL guardcolumn (Tosoh, 0008033), TSKgel GMPWXL column (Tosoh, 0008025) and a TSKgel G4000SWXL column (Tosoh, 0008542) attached in series. The columns and samples were kept at 30 °C to avoid gelation of the samples. Gelatin molecules were detected by a UV detector, at 214 nm. The average molecular weight was defined by measuring poly(styrene sulfonate) sodium salt standards (PSS Polymer Standards, PSS-pss kit) of various molecular weights, ranging from 891 Da to 976000 Da, together with the same benzoic acid internal standard. Using the WinGPC software package (PSS), an internal standard correction was performed and the average molecular weight and distribution were calculated using a calibration curve obtained from the standards.

The isoelectric point (IEP) was defined by completely demineralizing a 2% gelatin solution by adding an abundance of mixed bed ion exchange resin Amberlite IRN-150 (VWR, 40992.A9) to the gelatin. Conductivity was assessed using a conductivity meter (Consort, Multi-parameter analyzer C3010) with a platinum electrode (Pt 1000). After demineralizing the solution, the pH was measured, which is equal to the IEP, using a pHmeter (Mettler Toledo, Seven direct SD20).

For gel strength determination, gels were prepared by casting a 6.67% gelatin solution in a bloom glass mold (Brookfield, Type TA-GBB-2) with an inner diameter of 59 ± 1 mm and a height of 85 ± 2 mm. These gels were maintained at 10 °C for 17 h. The gel strength was measured using a texture analyzer (TA instruments, TA.XT plus C texture analyzer) with a 12.7 mm diameter plunger. The plunger was pushed into the surface of the gel over a distance of 4 mm and the force necessary for this (in grams) was the resulting gel strength of the gel.

The conductivity of the gelatin was measured in a 2% gelatin solution by using a Consort multi-parameter analyzer (C3010) with automatic temperature compensation and a platinum electrode (Pt 1000).

The ion content determination of sulfates and chlorides was performed using segmented flow analysis using a Technicon AutoAnalyzer II. Specific reactions were performed for each ion analyzed: Sulfates in aqueous solutions react with barium chloride (ChemLab, CL05.0214) at a pH of 2.5 to form the non-dissolvable barium sulfates. The unreacted barium reacts with methyl thymol blue (MTB) (ChemLab, CL00.1396) and from a blue chelate at a pH of 12.5. Barium chloride and MTB are equimolar and equivalent at the highest concentration of measurable sulfates. The amount of non-complexed, gray MTB, which can be measured at a wavelength of 460 nm, is therefore equal to the concentration of sulfates. Chlorides form a non-ionized mercury chloride when they react with mercury ions from a mercury thiocyanate (ChemLab, CL07.1121). This reaction releases a thiocyanate which can react with Fe^3+^ to form a complex with a red color, which can be measured at a wavelength of 480 nm, in equal concentration with the number of present chlorides. The determination of sodium was performed using a flame photometer (ISA Biologie Phf 106), where the sodium amount was calculated using the specific wavelength (588–589 nm) for sodium upon ignition. For this method, the gelatin was first removed from the solution by precipitating it with trichloroacetic acid (Merck, T4885). The content of sodium was calculated using a calibration curve made by measuring dilutions of a standard sodium solution (Merck, 1.70238). To eliminate the interference of the potassium signal, which has a wavelength upon ignition similar to sodium, potassium was added to the standard solution (Merck, 1.70230). The recovery of the measurement was also measured and taken into account while calculating the final amount of sodium.

The content of calcium was measured using a titration with EDTA (Merck, 324503) and calcone (Merck, 104594) as indicator in an alkaline environment. The indicator resulted in a color change from light red to light blue. The amount of calcium was calculated using following Equation 1:

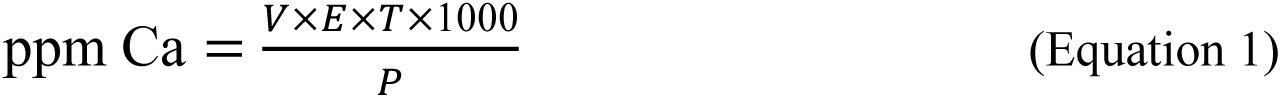

where V = volume titrated EDTA, E = equivalent mass (= 40.08), T = titer EDTA (=0.01), P = amount of gelatin (in grams).

For magnesium content, the amount of magnesium was measured using a titration with EDTA (Merck, 324503) and Eriochrome® Black T (Merck, 858390) as an indicator. This indicator has a wine red color in a aqueous solution with calcium and magnesium present at a pH of 10. With the addition of EDTA, the calcium and magnesium ions will form a complex and the indicator changes to a green-blue color. To calculate the total amount of magnesium, the amount of calcium calculated using Equation 1 was subtracted from the amount of complex formed (Equation 2).

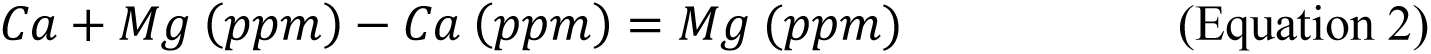

### Coacervation process to produce supporting baths by the CLADDING method

A detailed and theory-oriented protocol to generate a Crystal seLf-heAling embeDDing bIopriNtinG (CLADDING) support bath, using gelatin–gum arabic coacervation for bioprinting low-viscosity bioinks, with or without encapsulated cells, can be found in the **Supplementary methods.**

Briefly, after selecting the optimal gelatin, the ideal support bath was prepared by mixing 2% w gelatin type B from bovine bone 250 LBB8 (Rousselot Biomedical) with 0.5% gum arabic (G9752, Sigma-Aldrich) in 50% v of the total solvent (deionized water, dwater). The mixture were dissolved at 45 °C, vigorous stirring of 500 rpm, in a magnetic shaker (IKA, RH basic) for 1 h. After 1 h, the other 50% v of the total solvent (ethanol absolute, 100%) and 0.25% Pluronic F-127 (P2443, Sigma Aldrich) were added to the mixture. The pH was adjusted precisely to 5.8 using 1 M HCl. The solution was maintained at room temperature for 6 h, then at 4 °C for 12 h. The coacervates were distributed in 50 mL falcon tubes and centrifuged at 800 *g*, 5 min to remove ethanol remnants.

The hydrophobicity of the supporting bath was enhanced with either CaCl_2_ or Tween 20. For general use, washes with three times with 20 mM CaCl2, whereas if the transparency of the supporting bath is required, three times with 5% v/v Tween 20 in dwater. In each washing step with either CaCl_2_ or Tween 20, the coacervate was centrifuged (1000 *g,* 5 min) after which the supernatant was removed. The coacervates were mixed by shaking vigorously the falcon tube in the horizontal position, then the pellet was resuspended in another new washer solution. The third centrifugation, at 2000 *g* (5 min) for microparticle compaction resulted in an optimized supporting bath.

To measure gelatin–gum arabic microparticles, time-lapse microscopy was performed using a brightfield microscope (Eclipse Ti-E Nikon, Japan). Due to their electrostatic interaction, microparticles tended to aggregate immediately, forming a slurry that prevented their measurement. Thus, several videos were recorded beginning at the moment that microparticles were deposited on the microplate. Measurements of particle diameter before aggregation were performed using the imaging package of NIS Elements Software (Nikon).

### Analysis of the coacervation process in terms of yield as a function of pH, temperature, protein:polysaccharide ratio, and solvent

Aiming to achieve the optimized supporting bath, different parameters were evaluated. To obtain the highest level of precipitation, pH values of 4, 4.2, 4.4, 4.6, 4.8, 5.0, 5.2, 5.4, 5.6, 5.8, and 6.0 were investigated. To obtain the highest yield of coacervates, temperature, protein:polysaccharide ratio and solvent were investigated. Regarding temperature, maintaining at 25 °C or introducing a second drop of temperature to 4 °C were compared. As protein:polysaccharide ratio, 1:1 and 4:1 (gelatin:gum arabic) were analyzed. Regarding the solvent, 100% dwater for solubilization of gelatin:gum arabic, then 100% ethanol for precipitation versus a 50% ethanol/50% dwater solution since the beginning were investigated.

### Transparency of supporting baths

The macroscopic transparency of the support bath was assessed by placing them over text. Subsequently, light transittance was quantified at 400, 600 and 800 nm in a CLARIOstar plate reader (BMG Labtech). Dwater was used as baseline control and transmittance was calculated using Equation 3, according to a previously published protocol. ^[57]^

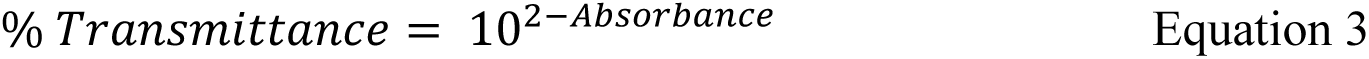

### Rheological characterization of supporting baths

Rheological characterization of the supporting baths was performed on a TA instrument DHR-2 Rheometer equipped with a Peltier temperature controller, with a cone-plate geometry (20 mm, 2° cone angle) and a 53 µm gap. Flow sweep curves were obtained between 0.01–100 s^-1^ shear rate (10 points/decade). Measurements were performed with steady-state sensing, with a maximum equilibrium time of 60 sec. Flow sweep curves were plotted as viscosity and stress vs. shear rate.

To analyze the self-healing properties of the supporting bath, a three-step flow assay was performed. The sample was submitted to a 0.1 s^-1^ shear rate for 2 min, followed by a 100 s^-1^ shear rate for 2 min, and then to a 0.1 s^-1^ shear rate for 2 min (1 point/s). These assays were plotted as viscosity vs. time. Amplitude sweeps were performed between 0.01–100 % oscillation strain, with a constant 1 Hz frequency. These assays were plotted as storage (G’) and loss (G’’) modulus vs. strain, and LVER was determined as the strain range with constant G’. Frequency sweeps were performed between 0.01–100 Hz oscillation frequency, with a constant 1% strain amplitude. These assays were plotted as storage (G’) and loss (G’’) modulus vs. frequency. G’ and G’’ were calculated within the plateau region.

### Formulation of BSMA and PLMA hydrogels and crosslinking strategy

The methacrylated platelet lysates (PLMA) used in this work were supplied by Metatissue, Portugal. They were obtained in a standardized manner and are commercially available (PLMA 100). The methacrylated bovine serum albumin (BSAMA) was obtained according to previously published protocols.^[58]^ Briefly, BSA (Sigma-Aldrich) was dissolved in carbonate bicarbonate buffer at 5% w at 37 °C for 1 h. After this, pH was adjusted to 9 with 5 M NaOH. While stirring, methacrylic anhydryde (Sigma-Aldrich) was added in a dropwise manner at a 10 v/v% to BSA. The solution was left to react at 500 rpm, 37 °C during 1h, after which pH was adjusted to 7.4. This solution was dialyzed against distilled water using a 3.5 KDa membrane for 72 h. The purified solution was filtered using 0.22 μm filters, frozen at -80 °C and freeze-dried for at least 7 days. The obtained powder (BSAMA) was stored at 4 °C and sealed with parafilm to prevent moisture.

Lyophilized PLMA and BSAMA were dissolved in modified Dulbecco’s phosphate-buffered saline (DPBS), without calcium chloride and magnesium chloride (D8537, Sigma-Aldrich) to a final concentration of 10% w/v for all experimets, except for the co-culture studies using hUVECs and hMSCs which was 30% w/v.. Stock solutions of 1 M sodium persulfate (SPS) photoinitiator and 0.1 M Ruthenium (Ru) photoinitiator (Photocrosslinking kit, 5248, Advanced BioMatrix) were dissolved separately in DPBS. SPS and Ru were mixed with the previously prepared solution of 10% or 30% PLMA/BSAMA in a 1:100 v/v ratio to achieve final concentrations of 0.01 M SPS and 0.001 M Ru. To analyze the photocrosslinking effect, solutions of 10% PLMA and BSAMA containing 0.001 M Ru/0.01 SPS photoinitiators were photocrosslinked using visible light at 455 nm and a distance of 1 mm from the LED source. Light intensities of 6, 11, 21 and 46 mW/cm^2^ and crosslinking time of 30, 45, 60 s were investigated. After optimization for the best crosslinking parameters determined by vial inversion test, 46 mW/cm^2^ and 60 s were defined to perform all subsequent experiments.

### Structural characterization of BSAMA and PLMA hydrogels

Proton nuclear magnetic resonance (^1^H NMR) spectroscopy was carried out for BSA, BSAMA, PL and PLMA by dissolving them at 10 mg.mL^−1^ in deuterium oxide (D_2_O, Merck) or in dimethyl sulfoxide-d_6_ (DMSO-d_6_). All the ^1^H NMR spectra were collected at room temperature using an Avance II 300 spectrometer (300.13 MHz). The deconvolution and simulation of NMR spectra were assessed through MestreNova® 9.0.1 software and the chemical shifts (*δ*) given in ppm. The degree of methacrylation was quantified by the fluoraldehyde o-Phthaldialdehyde (OPA) reagent solution (26025, ThermoFisher), which reacts with primary amines of proteins and amino acids enabling fluorescence detection and quantification. Samples were incubated in OPA (1:10) for 5 min and measured in a plate reader at excitation at 300–390 nm and emission at 436–475 nm.

Scanning electron microscopy (SEM) analysis was used to evaluate the structure of PLMA and BSAMA hydrogels and to calculate their porosity. Prior to observation, the hydrogels were freeze-dried for 24 h using a LyoQuest Plus Freeze-dryer (Telstar) and mounted on a stub. Then, samples were spin-coated with a thin layer of palladium-gold and images were captured using a Hitachi SU-3800 (Hitachi, Japan) scanning electron microscope with an acceleration voltage of 10 kV.

### Rheological characterization of PLMA and BSMA hydrogels

Rheological characterization of 10% w BSAMA and 10% w PLMA precursor solutions and crosslinked hydrogels was performed using a rheometer (Kinexus, Malvern), with an 8 mm parallel plate geometry and a 1 mm gap. To assess the viscosity of BSAMA and PLMA solutions, shear rate sweeps were conducted between 0.01–500 s^-1^ shear rate (5 points/decade), with steady-state sensing. These assays were plotted as viscosity vs. shear rate. To assess the linear viscoelastic region (LVR) of BSAMA and PLMA solutions, strain sweeps were performed between 0.1–100% oscillation strain, with a constant 1 Hz frequency. These assays are plotted as storage (G’) and loss (G’’) modulus vs. strain, and LVR was determined as the strain range with constant G’.

To evaluate the mechanical properties of BSAMA and PLMA crosslinked hydrogels, frequency sweeps were performed between 0.01–100 Hz, with a constant 0.1% strain. These assays were plotted as storage (G’) and loss (G’’) modulus vs. frequency. G’ and G’’ were determined for 1 Hz frequency. To characterize photocrosslinking, 10% (w/v) BSAMA and PLMA solutions in PBS were mixed with photoinitiators for obtaining final concentrations of 0.001 M Ru and 0.01 M SPS. The samples were submitted to an oscillation assay with a constant 0.1% strain and 1 Hz frequency. After two minutes, the solutions were irradiated with light (OmniCure S2000 Spot UV Curing System, 1 W/cm^2^, 320 –500 nm) for 13 min. These assays were plotted as storage modulus (G’) vs. time (t=0–7 min presented). Stabilization time was estimated as the instant when G’ varied less than 15%. G’ and G’’ were calculated before (average of values for t= 30–90 sec) and after (average of values for t= 840–900 sec) irradiation and compared to values obtained in frequency sweeps for the same conditions (1 Hz frequency, 0.1% strain).

### Stability, swelling index, and mass change of PLMA and BSAMA hydrogels over 50 days

To evaluate the stability and degradation of PLMA/BSAMA hydrogels, as well as the release of their components, droplets (20 µL) of PLMA and BSAMA were extruded, photocrosslinked in dried conditions, and soaked in MEM Alpha Medium with Gluta-MAX (32561-029, Gibco), supplemented with 10% (v/v) of fetal bovine serum (FBS, F7524, Sigma-Aldrich) for 50 days (37 °C). The masses of hydrogel droplets (hydrated PLMA/BSAMA) and lyophilized droplets (dried PLMA/BSAMA) were assessed on days 3, 10, 30, and 50 and compared to freshly prepared ones (day 0), as previously described.^[44]^ The supernatant was harvested on days 3, 10, 30, and 50 and analyzed using the Micro BCA protein assay (23235, ThermoFisher) for quantification of total protein released from PLMA/BSAMA hydrogels over time. The assay was performed following the protocol of the fabricant. Samples with culture media were used as a baseline control considering that culture media have protein components. Due to that, the Micro BCA protein assay was used instead of the standard BCA, as the signal has higher accuracy. Briefly, 150 µL of standard and samples were mixed with 150 µL of the working reagent (25:24:1, reagents MA, MB, MC) in a microplate and incubated for 2 h at 37 °C. A standard curve was prepared with the BSA solution provided by the kit. Absorbance was assessed in a plate reader at 562 nm and quantified relative to blank.

### Cell culture conditions

Human Mesenchymal Stem Cells (hMSC) at passage 5 (PT-2501, Lonza) were used. HMSCs were expanded by culturing 3000 cells/cm^2^ in T-225 flask until they reached 70% of confluence. Expansion basic medium composed by MEM Alpha Medium with GlutaMAX (32561-029, Gibco), supplemented with 10% (v/v) FBS (F7524, Sigma-Aldrich), was used. Media refreshments were performed every 2 days. Human Umbilical Vein Endothelial Cells (hUVECs) at passage 4 (C-12203, PromoCell) were used. HUVECs were expanded by culturing 5000 cells/cm^2^ in a T-175 flask until they reached 80% of confluence. Endothelial Cell Growth Medium 2 (EGM2, C-22011, PromoCell) was used. Media refreshments were performed every 2 days. All cell culture experiments were maintained at 37 °C, pH 7.2 under a 5% CO_2_ atmosphere.

### Bioprinting hMSCs in single and co-cultures with hUVECs in PLMA hydrogels using CLADDING

For bioprinting only hMSCs, the inks were prepared first by mixing 10% (w/v) PLMA in PBS or culture media with SPS and Ru photoinitiators (in a 1:100 v/v ratio to achieve final concentrations of 0.01 M SPS and 0.001 M Ru). The cell concentration of the bioink was 4 million hMSCs per mL of PLMA/Ru/SPS solution.

For bioprinting the co-culture of hMSCs and HUVECs, the inks were prepared following the same procedure but now containing 30% (w/v) PLMA in PBS instead of 10% (w/v). The cell concentration of the bioink was 10 million cells per mL of PLMA/Ru/SPS solution (co-culture of 2 million hMSCs and 8 million of hUVECs). Bioprinted constructs were cultured in EGM2 culture medium for 14 days, with media refreshments every 2 days.

A pressure-assisted extrusion bioprinting technique (BioScaffolder 3.1, Gesim – Gesellschaft für Silizium-Mikrosysteme mbH, Germany) was used to manufacture all constructs. The bioinks were loaded into a sterile standard syringe covered with aluminum foil and containing a sealing rubber positioned far from the solution to create a vacuum to avoid dripping due to their high fluidity (**Supplementary** Figure 23). The nozzle chosen had a long, straight, stainless-steel needle of 25 gauge, 0.25 mm of internal diameter, and 0.52 mm of external diameter (7018333, Nordson).

The optimal bioprinting parameters were 10 mm.s^-1^ for printing speed and 15–20 kPa for extrusion pressure. For photocrosslinking, a LED source with an intensity of 46 mW/cm^2^, at 455 nm for 60 seconds was employed. The photocrosslinking happened simultaneously with bioprinting, thus the LED source was coupled to the bioprinter to ensure reproducibility of light position.

To evaluate the extruded filaments of PLMA and BSAMA hydrogels in terms of size, resolution, and stability, optical microscopy pictures were taken and compared to the preset CAD model. All the measurements were performed either using the imaging package of NIS Elements Software (Nikon, Japan) or Image J.

### Biochemical assays and immunohistochemical analysis

To assess cell viability, live/dead assays were carried out with a staining solution consisting of 2.5 μM of ethidium homodimer-1 (EthD-1, E1169, Thermo Fisher) and 1 μM of calcein acetoxymethyl (Calcein AM, c3099, Thermo Fisher). The samples were analyzed with a fluorescence microscope (Eclipse Ti-E Nikon, Japan).

To analyze the metabolic activity of cells and bioprinted microtissues, a Presto Blue cell viability reagent (A13261, Life Technologies) was used. Samples were washed three times with PBS, followed by incubation with a presto blue solution in culture medium (1:9) for 3 h. Fluorescence was assessed at 560 nm for excitation and 590 nm for emission. Depending on the objective of the experiment, lower or higher cell concentrations were employed. Aiming to analyze cell cytotoxicity and reproducibility between batches of PLMA, a cell concentration of 0.5 million cells/mL was employed (96 well plate, 50.000 cells/well). Metabolic activity was assessed after 24 h, in triplicate. For selecting PBS or medium culture as the best solvent for preparing the bioinks, a cell concentration of 2 million cells/mL was employed (96 wellplate, 200.000 cells per well). Metabolic activity was assessed after 24 h and 7 days, in triplicate. For bioprinting PLMA/hMSCs bioinks, a cell concentration of 4 million cells/mL was used. After 24 h, 4 d, and 7 d, metabolic activity was assessed. For bioprinting trials, six replicates for each time point were assessed.

Immunohistochemical analysis was performed to determine the distribution of cells by staining cell nuclei with DAPI, cytoplasm with phalloidin and distinguishing hUVECs using the endothelial cell-specific marker CD31. Permeabilization and blocking were performed simultaneously with a solution of 1% Trition-X-100, 5% goat serum, 0.05% Tween 20, and 1% bovine serum albumin (BSA) in PBS, for 24 h at 4 °C, under mild agitation. The samples were then incubated for 24 h at 4 °C, under mild agitation, with primary antibody solution containing CD31 antibody (M0823, Dako, 1:100), 0.1% Triton X-100, 5% goat serum, 0.05% Tween20, and 1% BSA in PBS. The samples were then washed with a wash buffer composed of 0.05% Tween 20 and 1% BSA in PBS, and left for 24 h at 4 °C, under mild agitation, to remove unbound antibodies. Secondary antibody solution of goat anti-mouse conjugated with Alexa Fluor 647 (Thermo Fisher Scientific) were prepared in wash buffer and incubated for 24 h at 4 °C, under mild agitation. Bioprinted constructs were incubated with Alexa Fluor 568-phalloidin (Thermo Fisher Scientific, 1:100) for 90 min at room temperature (RT), washed 3 times with PBS, and with phalloidin (1:100) for 90 min, washed 3 times with PBS, and incubated with DAPI (1:100) for 1 h at RT. After 3 washing steps with PBS for 5 min each, bioprinted contructs were analyzed with a fluorescence microscope (Eclipse Ti-E Nikon, Japan) or Confocal Laser Scanning Microscope (Leica TCS SP8, Germany). The images from the same group of analyses were taken on the same day, in triplicate, using the same settings.

## Statistical analysis

Statistical analysis was conducted using Prism software (8.4.3 version, GraphPad). One-way or two-way ANOVA was employed with Tukey’s *post hoc* comparison to evaluate statistical significance. All assays herein presented were performed at least in three technical replicates. The data are expressed as mean ± standard deviation (SD) values or represented using standard deviation bars in the graphs.

## Acknowledgements

The authors acknowledge the support of the European Union’s Horizon 2020 research and innovation program under grant agreement No 953169 (**InterLynk**). Helena P. Ferreira and Inês Gonçalves acknowledge the financial support from Portuguese FCT for PhD grant 2020.04712.BD and contract CECINST/00091/2018/CP1500/CT0013. Rita Sobreiro-Almeida and Catarina Custódio acknowledge the Portuguese Foundation for Science and Technology for their individual grant 2022.04605.CEECIND and 2020.01647.CEECIND, respectively. The authors would also like to thank Thomas Van Gansbeke (Rousselot Biomedical) for support in the characterization of molecular weight and ion content of the gelatins used for this study.

Appendix A. Supplementary data

