## Supplementary Information for "Embedding bioprinting of low viscous, photopolymerizable blood-based bioinks in a self-healing transparent supporting bath"

### Supplementary methods

#### Coacervation process to produce supporting baths

Due to the conceptual character of this study, we are offering a detailed and theory-oriented protocol for generating a Crystal seLf-heAling embeDDing bIopriNtinG (CLADDING) support bath, using gelatin–gum arabic coacervation for bioprinting low-viscosity bioinks, with or without encapsulated cells,

**Coacervate preparation:**

a) Mix gelatin to gum arabic (4:1) in deionized water (dwater) at 45 °C corresponding of 50% of the total solvent. Example of a 1 L solution: 20 g of gelatin, 5 g of Arabic gum in 500 mL of dwater (add in small portions, not all at once to avoid formation of clumps). For our supporting bath, gelatin type B from bovine bone (type II called herein) 250 LBB8 (Rousselot), and gum arabic (G9752, Sigma-Aldrich) were used.
b) Perform vigorous stirring (500 rpm, IKA RH basic) in a magnetic shaker at 45 °C, for at least 1 hour. Use a thermometer for assuring that the solution is at 45 °C.
c) After 1 hour, a complete dissolution of gelatin and gum arabic should be seen. The transmittance values of these procedure can be seen in Figure 2E.

**Micro-coacervation process**

d) Add the other 50% of the solvent of ethanol absolute (100%) and Pluronic F-127 (2.5 g/L). For the optimized support, Pluronic® F-127 (P2443, Sigma Aldrich) was used.
e) Right after, check the pH. It should be around 6.2-6.6 and the solution is transparent. Adjust the pH precisely for 5.8 using 1 M HCl and the solution will become turbid. The transmittance values of these procedure can be seen in Figure 2E. This plays a relevant role in the emulsion formation since the decrease of pH is one of the coacervate agents.
f) Turn off the heating platform and keep stirring for around 15 min.
g) It will take some time for decreasing the temperature of the system and the gelatin-gum arabic microparticles to precipitate. Close the flask with a lid to avoid ethanol evaporation and place it at room temperature for 6 hours.

**Macro-coacervation process**

h) After the first temperature decrease, place the flask at 4 °C overnight. After 12 hours, the coacervate will be formed on the bottom of the flask. These double step of decreasing temperature acts as the other coacervate agent.

i) Using pipettes, remove the liquid equilibrium phase located above the coacervate, avoiding disturbance on the coacervate. By our experience, for each 1L of a solution prepared, around 600 mL of equilibrium phase should be removed.

j) Distribute the coacervate in 50 mL falcon tubes to have at least half of coacervate and half of fluid phase on each.

k) Centrifuge the tubes at 800 g, 5 min to remove ethanol remnants. After centrifugation, a clear phase separation can be seen. Do not add water for washing at this step.

**Washing steps to increase hydrophobicity or obtain transparency**

l) For optimized bioprinting of water-soluble bioinks, the hydrophobicity of the supporting bath should be enhanced. We recommend the use of $CaCl_2$ or Tween 20. For general use, three times with $CaCl_2$ 20 mM, whereas if the transparency of the supporting bath is required, three times with 5% v/v Tween 20 in dwater. In each washing step with both $CaCl_2$ or Tween 20, the coacervate is centrifuged (1000 g, 5 min) and the supernatant is removed. The coacervates are mixed vigorously by shaking vigorously the falcon tube in the horizontal position and the pellet is resuspended in another new washer solution. Do it slowly since the $CaCl_2$ or Tween 20 solutions should act as a washing agent on the microparticles. On the third centrifugation, we recommend 2000 g (5 min) for microparticle compaction, which results in an optimized supporting bath.

**Use and Storage**

m) Using a spatula or gel pipettes, the supporting bath can be spread on wellplates, flasks, or petri dishes, right before bioprinting. If it will not be used immediately, we recommend storing the supporting bath on step **j**. On this condition, the coacervates stay hydrated by the equilibrium phase and can be maintained at 4 °C for at least two months.

**Support bath removal**

n) Since it is a coacervation between gelatin-arabic gum, the arabic gum is not realized immediately after temperature increase such as gelatin. Thus, after bioprinting, add media solution on top of the scaffold/construct with the support bath and place the plate in the cell incubator (37 °C). The gelatin will melt after 30 min and the support will become liquid. Go gradually replacing media culture over some days to remove the support gradually while avoid breaking your printed constructs.

### Supplementary figures

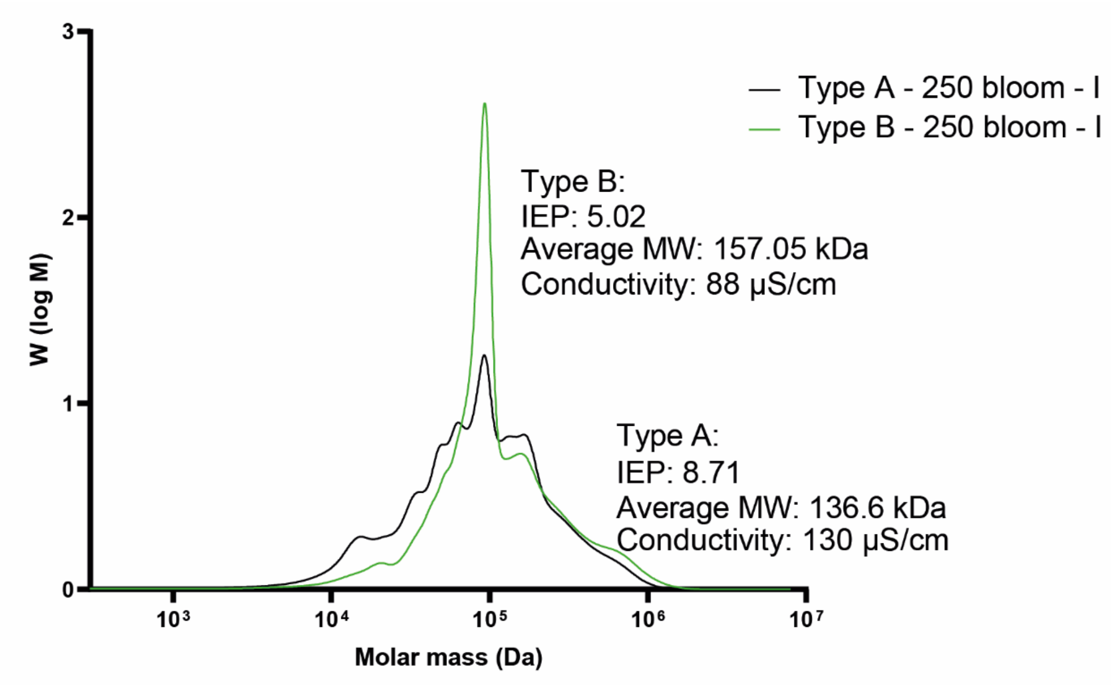


**Supplementary Figure 1:** Comparing the properties of gelatins type A and B with defined bloom of 250.

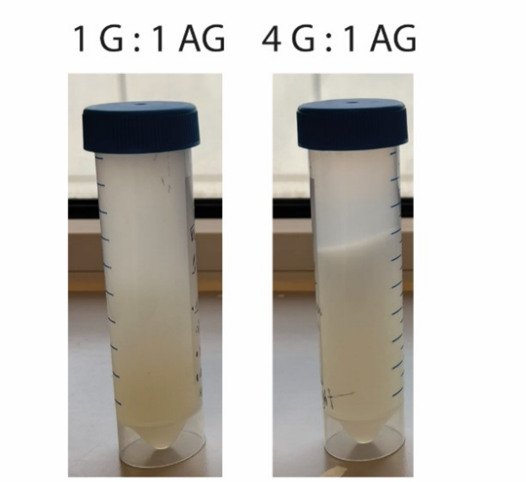


**Supplementary Figure 2:** Precipitation step using two ratios of gelatin and gum arabic (1:1 and 4:1). Using 1:1, it is not possible to obtain a biphasic system and the equilibrium phase is mixed into the coacervate.

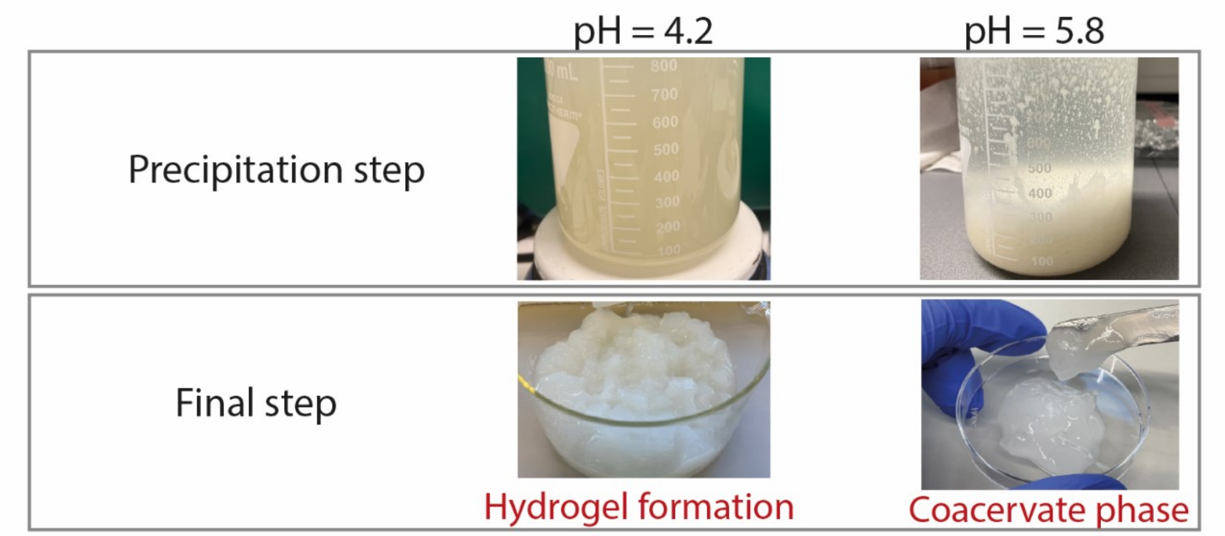


**Supplementary Figure 3:** Investigation of the optimum pH during macro-coacervation step of gelatin and gum arabic. At 4.2, a hydrogel was obtained in the final step. At 5.8, a proper coacervate was obtained.

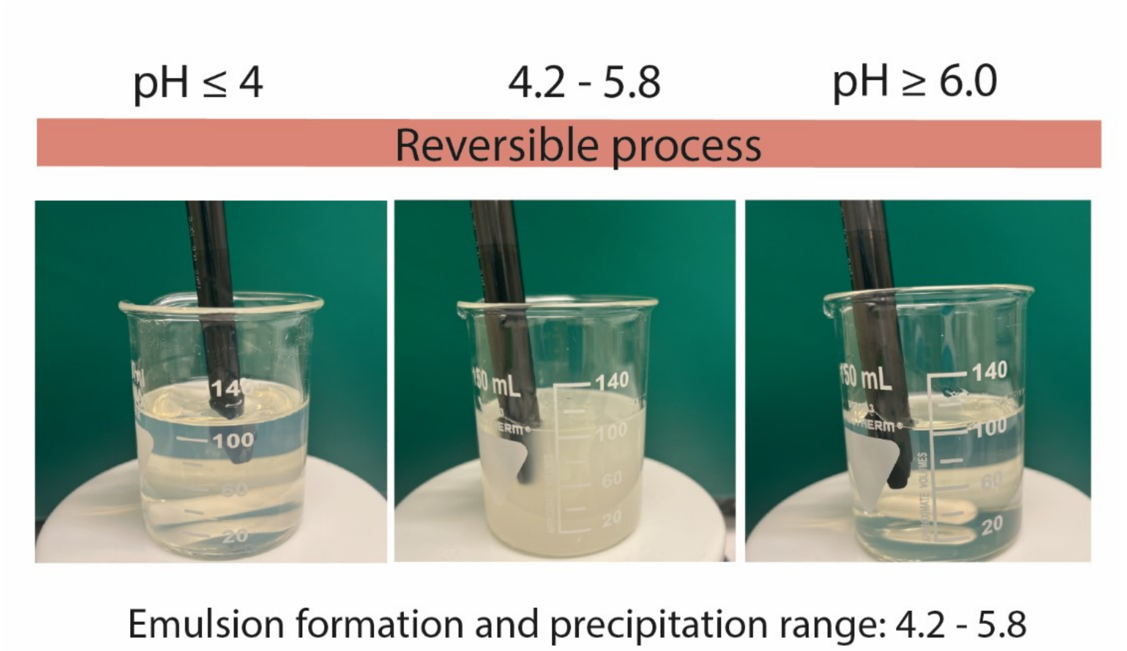


**Supplementary Figure 4:** In the macro-coacervation step, pH can be altered from 4 until 6 in a reversible process. Emulsion formation happens only from 4.2 – 5.8, achieving high yield at 5.8.

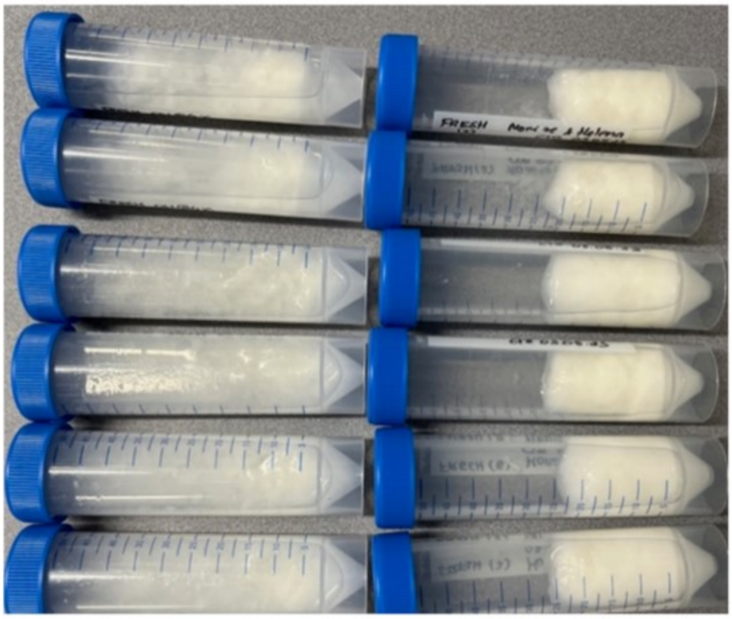

**Supplementary Figure 5:** In the macro-coacervation step, the second temperature drop to 4 °C enhanced the compactness of the coacervate (right) when compared to 6 h maintained at room temperature (left). The micro and macro coacervation is performed in flasks. The pictures were taken after concluding the whole process in both conditions (one temperature drop to 25 ºC in the left image and two temperature drops to 25 ºC, then to 4 ºC, in the right image).

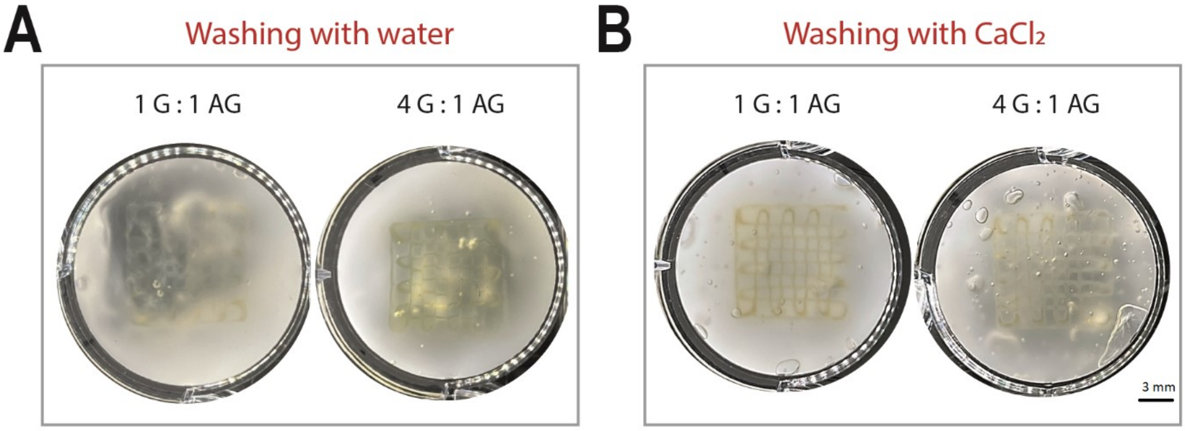


**Supplementary Figure 6:** Printability of PLMA in supporting baths using two ratios (1:1 and 4:1), washed with water and with $CaCl_2$. Since PLMA is a water-based bioink, it clearly spreads in the most hydrophilic supporting bath and well-defined filaments were not obtained (scale bar = 3 mm).

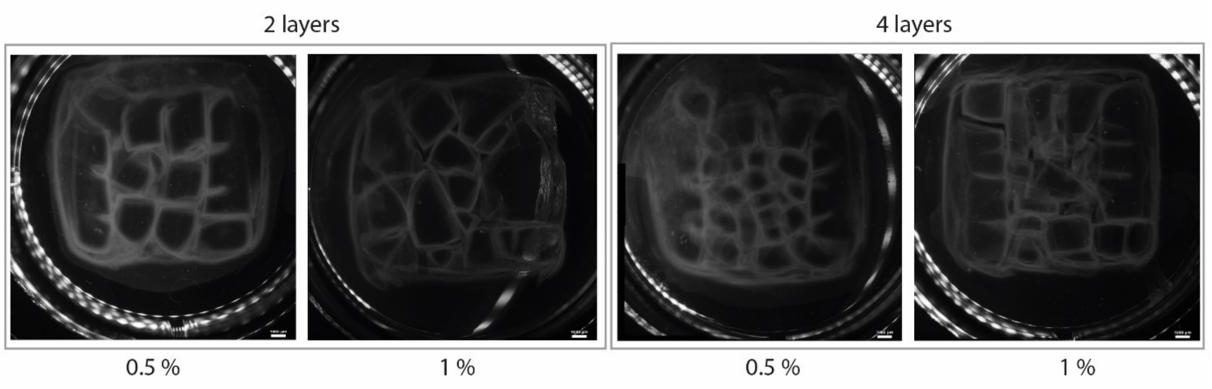


**Supplementary Figure 7**: Support bath made of a full block of gelatin, 0.5 and 1 % (w/v). When bioprinting inside these supports, scaffolds with squared geometry were obtained showing that the needle was facing a high resistance to move. After bioprinting, pieces of gelatin could be seen, which shows that the support bath was stiffer that should be and did not presented the envisioned self-healing behavior (scale bar = 1 mm).

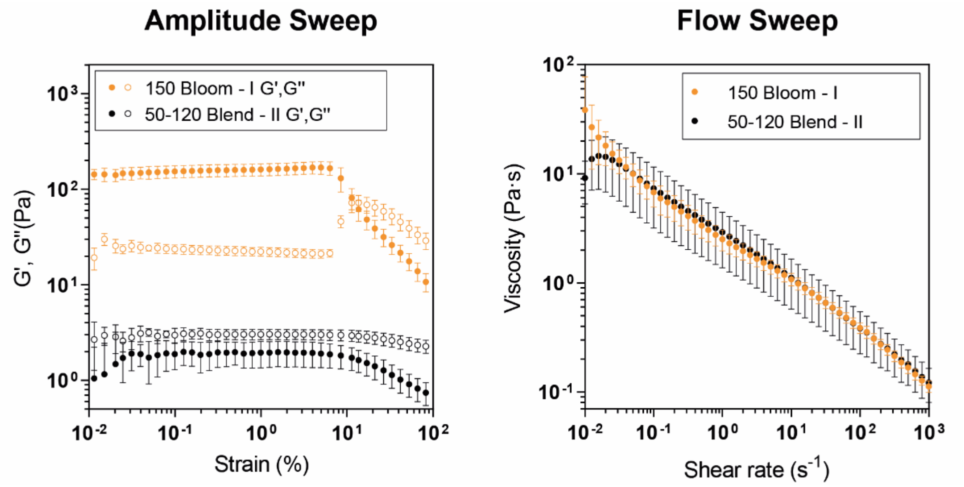


**Supplementary Figure 8**: Amplitude and flow sweep of gelatins with defined bloom of 150 and non-defined bloom of 50-120 blend gelatin.

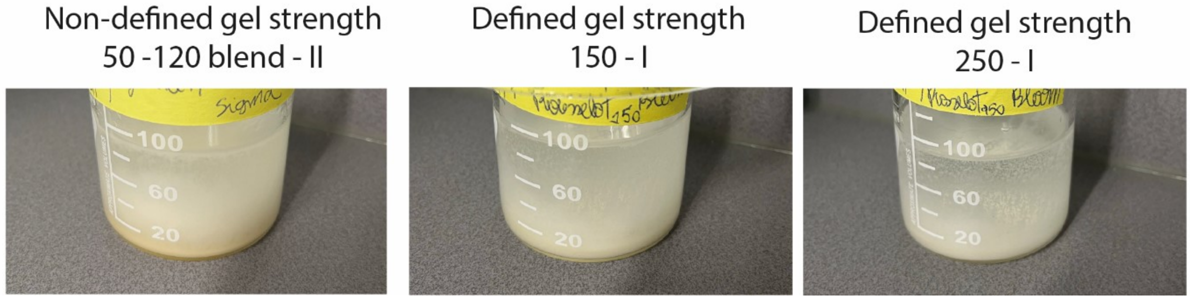


**Supplementary Figure 9:** Visually, precipitation happens similarly when using both defined and non-defined gel strength, which emphasized the need of proper characterization of the chosen gelatin.

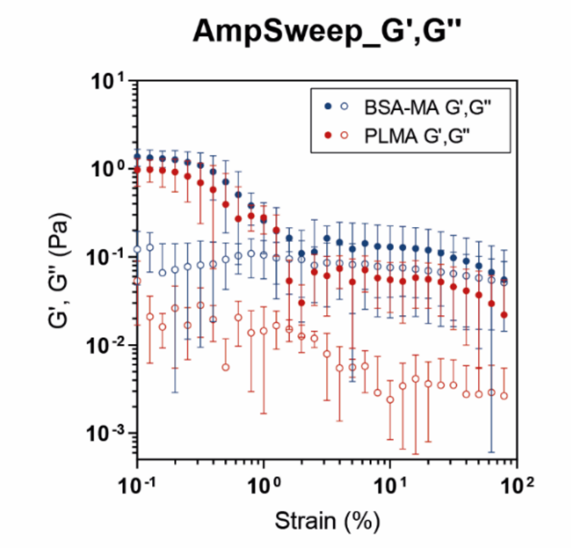


**Supplementary Figure 10:** Amplitude sweep of BSAMA and PLAMA bioinks.

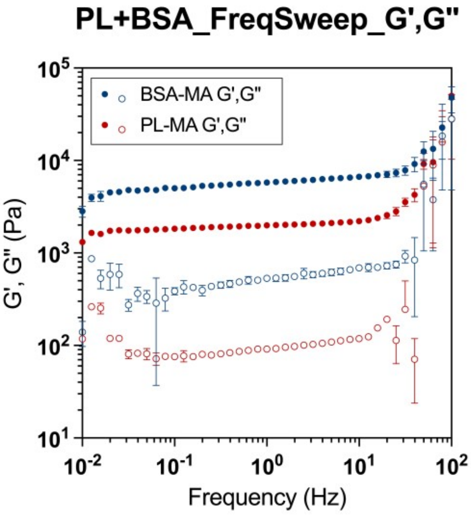


**Supplementary Figure 11:** Frequency sweeps of BSAMA and PLAMA hydrogels.

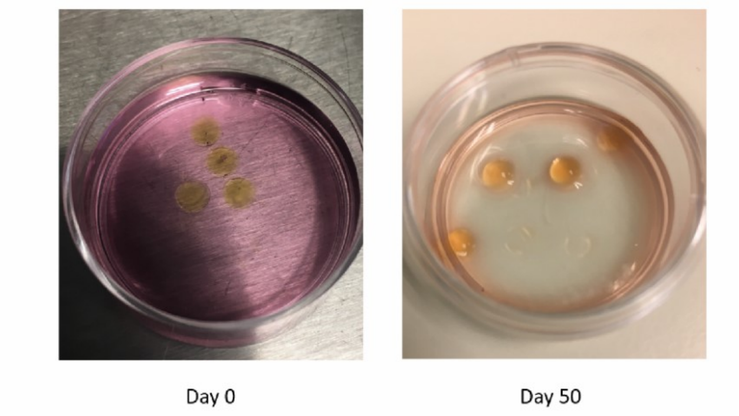


**Supplementary Figure 12:** Stability of BSAMA up to 50 days in culture (100 ± 8 % at day 1 and 99.2 ± 4 % at day 50).

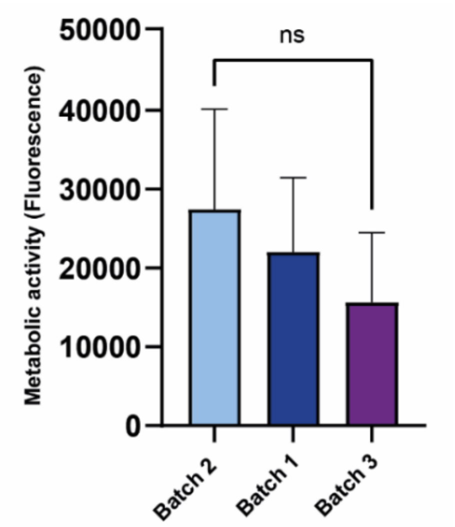


**Supplementary Figure 13:** Comparison of metabolic activity of hMSCs after 24 h when encapsulated in three independent batches of PLMA.

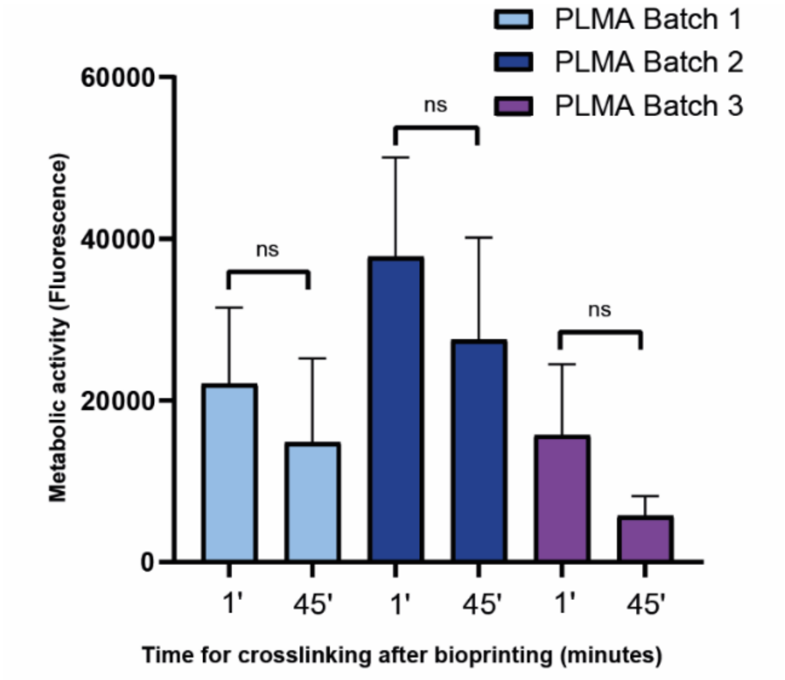


**Supplementary Figure 14:** Metabolic activity of hMSCs when maintained with methacrylated PL solution for 1 minute or 45 minutes before photocrosslinking. From a technical point of view, 45 minutes is enough time for loading this liquid bioink in the cartridge and performing all the necessary calibrations of the bioprinter prior bioprinting and photocrosslinking.

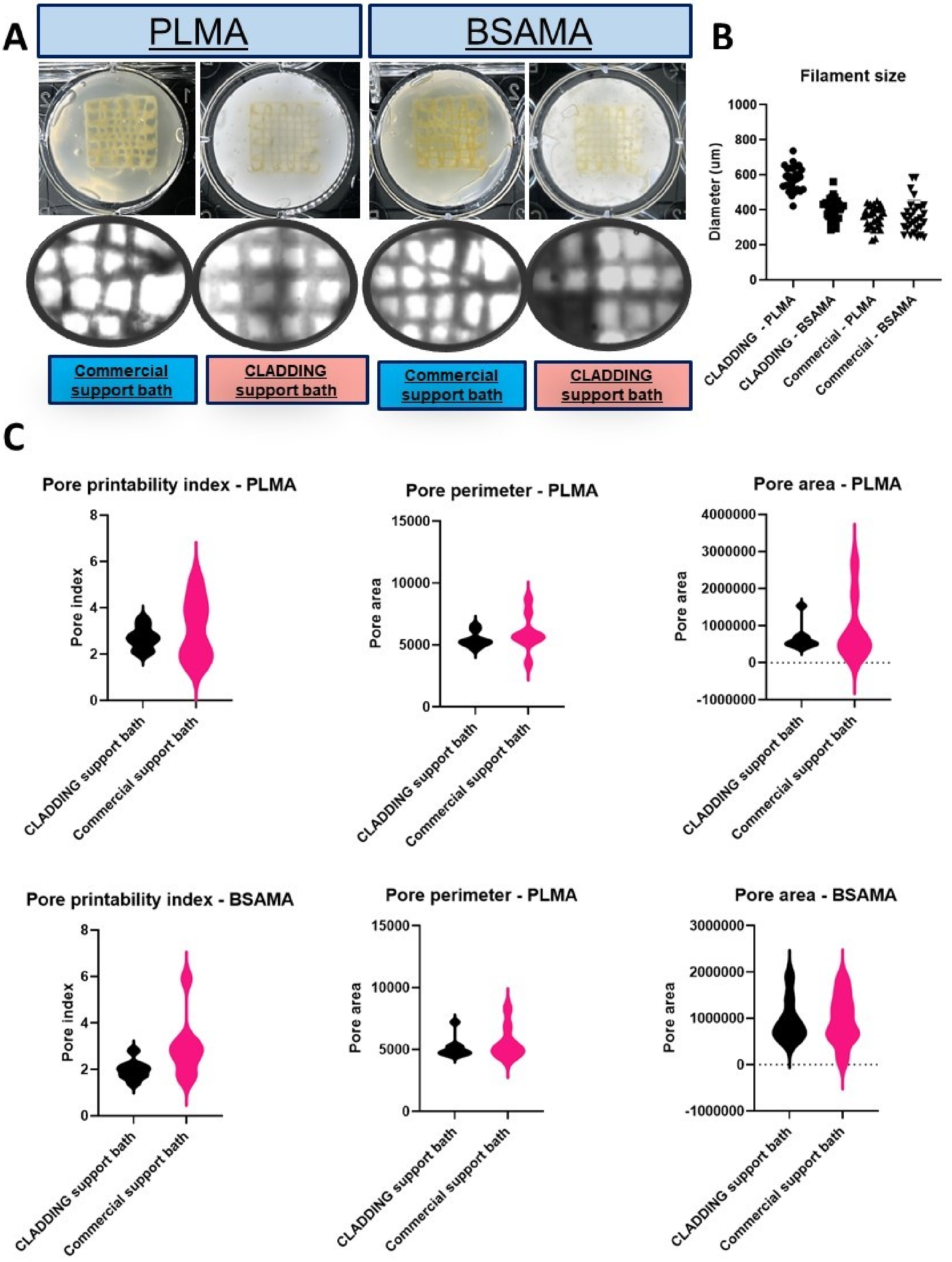


**Supplementary Figure 15:** Comparing our support bath with the commonly used commercial one (LifeSupport, FluidForm$^{TM}$) in terms of **A)** printing fidelity, **B)** filament size and **C)** pore parameters (printability index, area and perimeter).

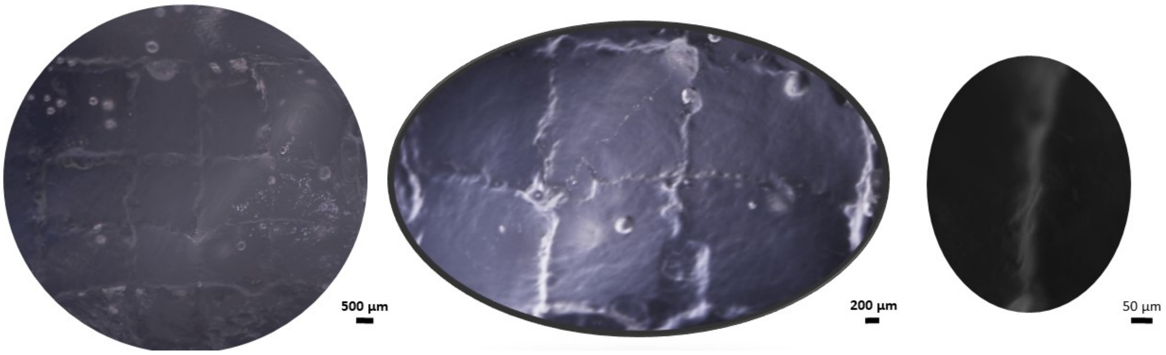


**Supplementary Figure 16:** Challenging the system with a much smaller diameter as technically possible with current available needles for embedding bioprinting (310 µm outer diameter, 30 gauge/150 µm of internal diameter), filaments of 142 ± 36 µm with PLMA were obtained in a intact structure, though with less resolution and stability than using the 25 gauge. Scale bar from the left to the right are 500, 200 and 50 µm.

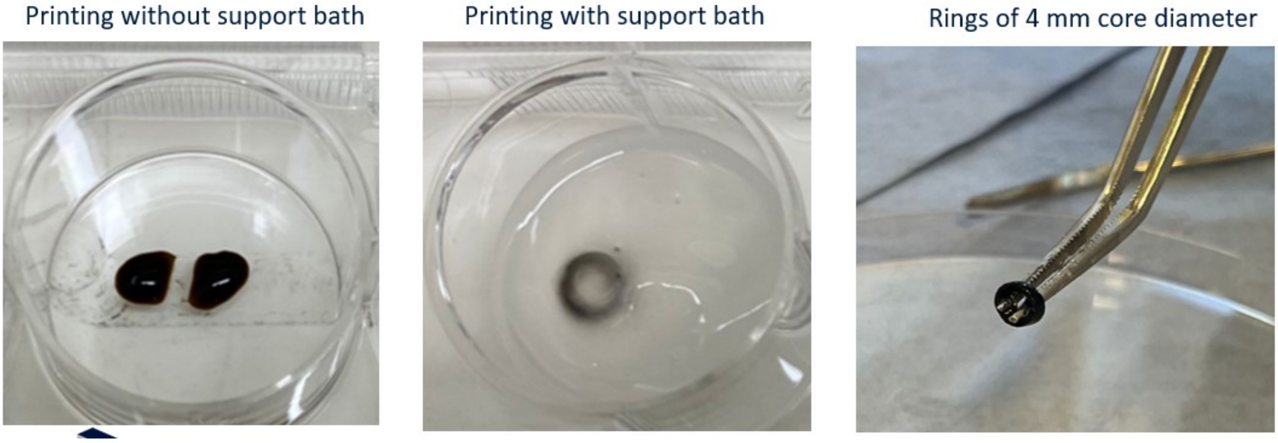


**Supplementary Figure 17:** Printing PEG-based bioinks using CLADDING shows the adaptability of this approach to other highly fluid low-viscous bioink synthetic formulations that are otherwise impossible to print.

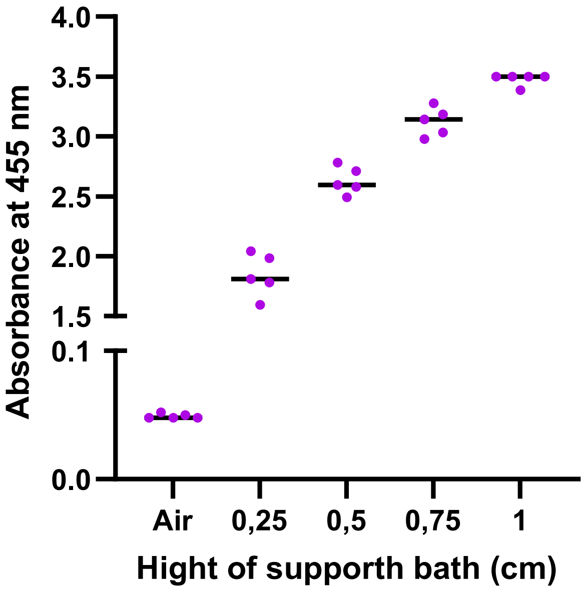


**Supplementary Figure 18:** Absorbance of 455 nm blue light by the support bath depending on the high of the bath that implies in the depth of the light.

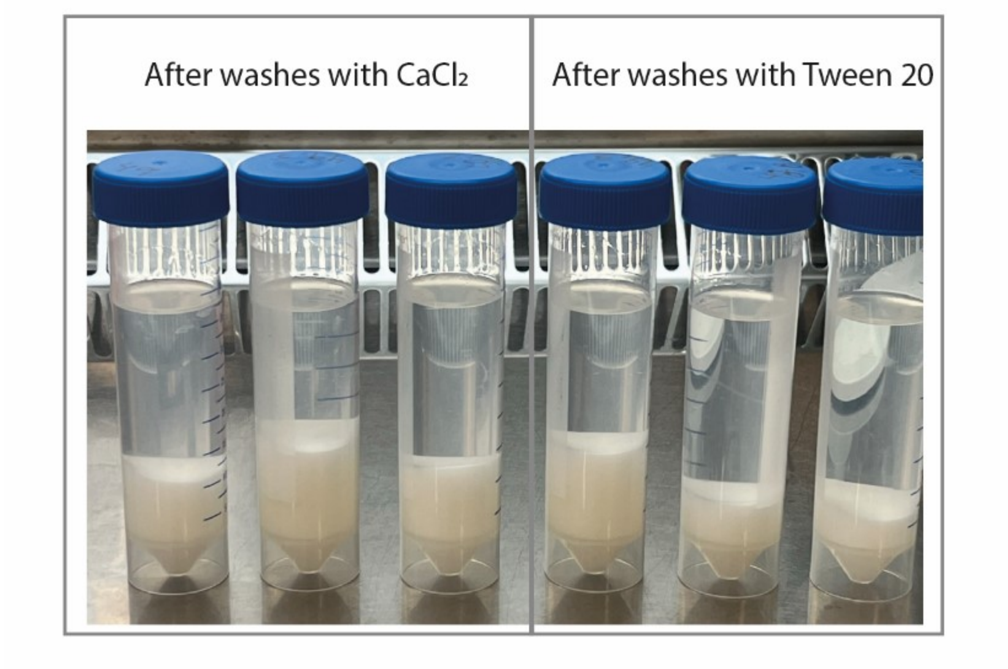


**Supplementary Figure 19:** Well-defined biphasic system can be obtained after washing steps both 5 % Tween 20 and 20 mM $CaCl_2$.

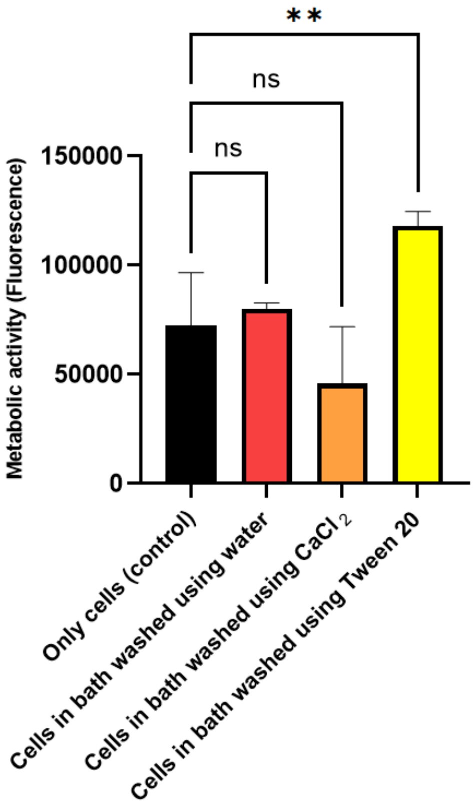


**Supplementary Figure 20:** Metabolic activity of hMSC after 24 h maintained in monolayer (control, without supporting bath), in supporting bath washed with water and in both 5 % Tween 20 and 20 mM $CaCl_2$.

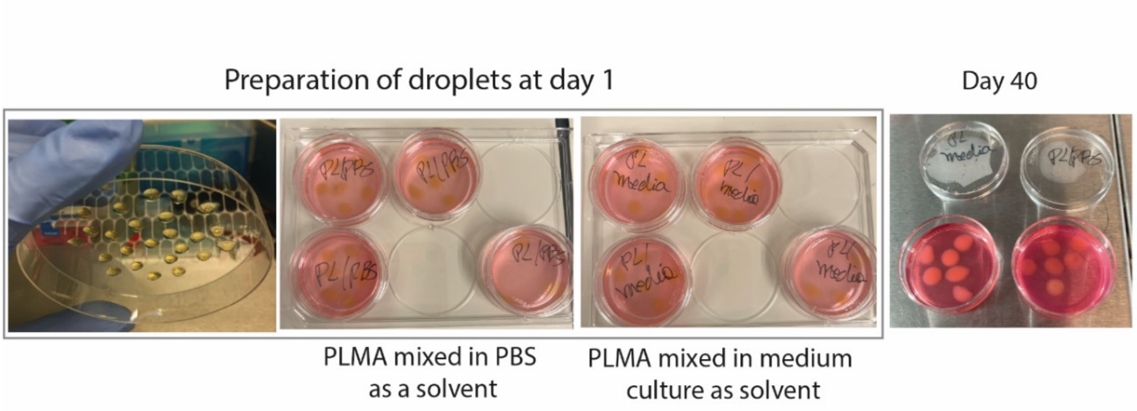


**Supplementary Figure 21:** Droplets of PLMA made using culture medium or PBS as solvent. These PLMA/medium culture or PLMA/PBS were photocrosslinked and manintained in cell culture conditions over 40 days.

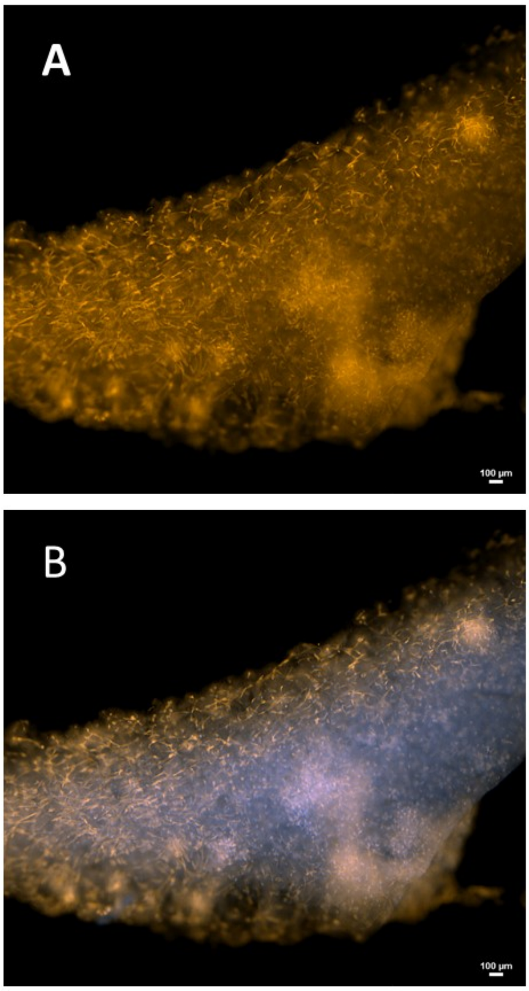


**Supplementary Figure 22**: **A)** Bioprinted filament stained with phalloidin (orange) to emphasize only the f-actin distribution of the cells. In spots that there are no cell aggregation or spheroids, the cells are very elongated and aligned. **B)** Same bioprinted filament merged with cell nuclei stained (DAPI, blue). Scale Bar: 100 um.

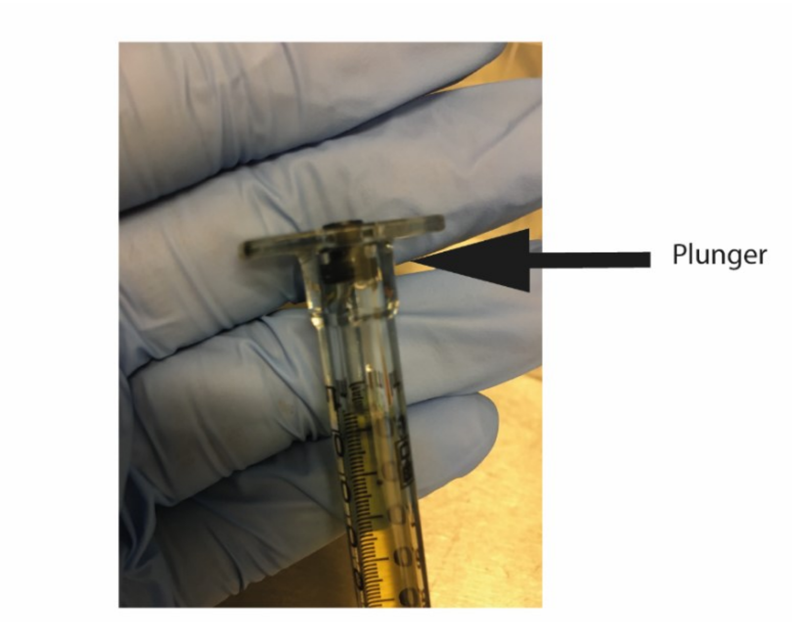


**Supplementary Figure 23:** Standard syringe containing a sealing rubber positioned far from the solution to create a vacuum to avoid dripping the bioink due to their high fluidity.

### Supplementary videos

**Supplementary Video 1:** Stable rings fully composed of PLMA were obtained when bioprinting multiple passes of PLMA (10 layers), along with simultaneously photocrosslinking.

**Supplementary Video 2:** Self-healing behavior of CLADDING support bath after washes with $CaCl_2$, Tween 20 and dwater (as a control).

**Supplementary Video 3:** Bioprinting 10 layers of PLMA bioink moving in Z, aiming to print a stable cylinder.

**Supplementary Video 4:** Repetitive cycles of placing and taking out porous constructs after support bath removal. Constructs were fully manufactured using PLMA and high concentration of hUVECs and hMSCs.

### Supplementary Tables

**Supplementary Table 1:** Additives combined with PL and PRP for enabling printing or bioprinting.

| | **Additives for increasing viscosity** | **Crosslinking strategy** | **Bioprinting system** | **Authors** |
|---|---|---|---|---|
| **Platelet lysates (PL)** | None | Photo - Visible light Ru/SPS | Extrusion with a conventional syringe and enhanced supporting bath | Present study |
| | Gelatin-alginate | Photo UV Lap | Extrusion | Min et al., 2021 [9] |
| | Gelatin methacryloyl (GelMA) | | | Daikuara et al., 2021 [10] |
| | Cellulose nanocrystals (CNC) | Ionic $Ca^{2+}$, thrombin, aldehyde-CNC | Extrusion with a double-barrel and supporting bath | Mendes et al 2020 [11] |
| | Alginate | Ionic $Ca^{2}$ | Extrusion | Faramarzi et al 2018 [12] |
| | Microgels (Norbornene-hyaluronic acid) | Photo UV Lap | | Mendes et al., 2021 [13] |
| **Platelet-rich plasma (PRP)** | Gelatin-alginate | Ionic $Ca^{2}$ | Extrusion in-situ bioprinting of a rat dorsal wound | Zhao et al, 2022 [4] |
| | Silk fibroin | Physical Beta-sheet formation | Extrusion | Li et al., 2020 [14] |
| | Extracellular matrix (ECM)-GelMA | Photo UV LAP | | Zhu et al., 2023 [15] |
| | GelMA | Photo UV Irgacure | | Irmak et al., 2021 [16] |
| | | Photo UV LAP | | Ibanez et al., 2022 [17] |
| | Alginate-agarose | Ionic $Ca^{2+}$/thrombin | | Zou et al., 2020 [18] |

**Supplementary Table 2:** Comparing gelatin-gum arabic ratios and solvents during micro-coacervate phase in terms of yield and waste**.** For each test, a 100 mL solution was prepared in three independently replicates.

**Micro-coacervate phase**

| **Parameters** | Liquid phase (waste, in mL) | Highly compacted coacervate (waste, in grams) | Average of waste (grams) | Fluid coacervate (collected, in grams) | Embedding support after centrifugstion of the collected fluid coacervate (ready to use) | Average of embedding support obtained (grams) |
|---|---|---|---|---|---|---|
| Macromolecule ratio: 1 gelatin : 1 gum arabic<br>Solvent: 100% water for solubilization<br>100% ethanol for precipitation | 69.0 | 12.6 | **12. 8 ± 0.6** | 12.2 | 6.0 | **6.1 ± 0.4** |
| | 68.0 | 12.3 | | 10.4 | 6.5 | |
| | 68.5 | 13.4 | | 8.8 | 5.8 | |
| Macromolecule ratio: 4 gelatin : 1 arabic gum<br>Solvent: 100% water for solubilization<br>100% ethanol for precipitation | 54.7 | 7.8 | **7.8 ± 0.1** | 23.4 | 12.7 | **12.5 ± 0.2** |
| | 52.7 | 7.9 | | 26.5 | 12.3 | |
| | 49.0 | 7.8 | | 29.9 | 12.5 | |
| Macromolecule ratio: 4 gelatin : 1 arabic gum<br>Solvent: 50% water 50% ethanol mixture for solubilization | 25 | 8.2 | **9.5 ± 1.2** | 47.5 | 5.9 | **6.3 ± 0.5** |
| | 16 | 10.5 | | 51.0 | 6.8 | |
| | 18 | 9.8 | | 51.8 | 6.3 | |

**Supplementary Table 3**: Comparing steps of temperature reduction during macro-coacervate phase in terms of yield and waste. For each test, a 100 mL solution was prepared in three independently replicates.

**Macro-coacervate phase**

| **Parameters** | Liquid phase (waste, in mL) | Highly compacted coacervate (waste, in grams) | Average of waste (grams) | Fluid coacervate (collected, in grams) | Embedding support after centrifugstion of the collected fluid coacervate (ready to use) | Average of embedding support produced (grams) |
|---|---|---|---|---|---|---|
| Macromolecule ratio: 4 gelatin : 1 arabic gum Solvent: 100% water for solubilization 100% ethanol for precipitation<br><br>Two steps of temperature reduction:<br>- 45 to 25 °C (6 hours)<br>- 25 to 4 °C (12 hours) | 54.7 | 7.8 | **7.8 ± 0.1** | 23.4 | 12.7 | **12.5 ± 0.2** |
| | 52.7 | 7.9 | | 26.5 | 12.3 | |
| | 49.0 | 7.8 | | 29.9 | 12.5 | |
| Macromolecule ratio: 4 gelatin : 1 arabic gum Solvent: 100% water for solubilization 100% ethanol for precipitation<br><br>One step of temperature reduction:<br>- 45 to 25 °C (6 hours) | 76.5 | 7.7 | **7.7 ± 0.2** | 7.0 | 1.8 | **1.8 ± 0.3** |
| | 76.7 | 7.8 | | 6.0 | 1.5 | |
| | 78 | 7.5 | | 7.5 | 2.0 | |
